# DNA and RNA-SIP reveal *Nitrospira spp.* as key drivers of nitrification in groundwater-fed biofilters

**DOI:** 10.1101/703868

**Authors:** Arda Gülay, Jane Fowler, Karolina Tatari, Bo Thamdrup, Hans-Jørgen Albrechtsen, Waleed Abu Al-Soud, Søren J. Sørensen, Barth F. Smets

## Abstract

Nitrification, the oxidative process converting ammonia to nitrite and nitrate, is driven by microbes and plays a central role in the global nitrogen cycle. Our earlier metagenomics, *amoA*-amplicon, and *amoA*-qPCR based investigations of groundwater-fed biofilters indicated a consistently high abundance of comammox *Nitrospira*, and we hypothesized that these non-classical nitrifiers drive ammonia-N oxidation. Hence, we used DNA and RNA stable isotope probing (SIP) coupled with 16S rRNA amplicon sequencing to identify the active members in the biofilter community when subject to a continuous supply of NH_4_^+^ or NO_2_^−^ in the presence of ^13^C-HCO_3_^−^ (labelled) or ^12^C-HCO_3_^−^ (unlabelled). Allylthiourea (ATU) and sodium chlorate were added to inhibit autotrophic ammonia- and nitrite-oxidizing bacteria, respectively. Our results confirmed that lineage II *Nitrospira* dominated ammonium oxidation in the biofilter community. A total of 78 (8 in RNA-SIP and 70 in DNA-SIP) and 96 (25 in RNA-SIP and 71 in DNA-SIP) *Nitrospira* phylotypes (at 99% 16S rRNA sequence similarity) were identified as complete ammonia- and nitrite-oxidizing, respectively. We also detected significant HCO_3_^−^ uptake by *Acidobacteria subgroup10, Pedomicrobium, Rhizobactera*, and *Acidovorax* under conditions that favoured ammonium oxidation. Canonical *Nitrospira* alone drove nitrite oxidation in the biofilter community, and activity of archaeal ammonia oxidizing taxa was not detected in the SIP fractions. This study provides the first *in-situ* evidence of ammonia oxidation by comammox *Nitrospira* in an ecologically relevant complex microbiome.

## Introduction

Nitrification, the stepwise oxidation of ammonia (NH_3_) to nitrite (NO_2_^−^) and nitrate (NO_3_^−^), supplies the substrates for processes that initiate the loss of reactive nitrogen from the biosphere as N_2_. Understanding the organisms and environmental controls that drive nitrification is important as it controls global homeostasis of the N cycle. In engineered environments, complete nitrification is often desired: this is essential when waters are prepared and distributed for human consumption. Residual NH_3_ or NO_2_^−^ - the result of incomplete nitrification - renders the water biologically unstable and unsafe for human consumption. Hence, biological systems for source water treatment are contingent on nitrifying prokaryotes. Based on evolutionarily conserved taxonomic (small subunit, 16S rRNA) and functional (e.g. ammonia monooxygenase, *amoA*) gene surveys, *Nitrosomonas* (1–4), *Nitrosoarchaeum*, and *Nitrososphaera* have been identified as the abundant ammonium oxidizing prokaryotes (AOP) and *Nitrospira* (5, 6) as the abundant nitrite oxidizing prokaryotes (NOP) in drinking water treatment systems, consistent with the classical assumption of division of labor in the two nitrification steps.

Our previous studies on rapid gravity sand filters (RGSF), used in potable water preparation from groundwater, revealed nitrifying microbial communities with *Nitrospira* far more abundant than *Nitrosomonas* (7),with several *Nitrospira* genomes containing genes for ammonia oxidation (8), and with an abundance of comammox (complete ammonia oxidizing) *amoA* over AOB *amoA* genes (9). Together with the concurrent discovery of comammox *Nitrospira* strains by others (10–12), this suggested that comammox *Nitrospira* may drive ammonia oxidation in the examined groundwater-fed RGSFs. In addition, like *Nitrospira*, several *Acidobacterial*, and γ- and α-*proteobacterial* taxa were at consistently higher abundance than *Nitrosomonas* questioning their potential role in nitrification, as NH_3_ is the primary growth substrate entering the filters (7, 13–15). Identifying the active ammonia and nitrite oxidizing organisms is essential not only for making predictions for engineering purposes, but also for understanding the niches and biodiversity of nitrifiers. There has been a rapidly increasing documentation of global comammox *Nitrospira* occurrence across a myriad of habitats ranging from the subsurface, to soils to sediments, and from groundwaters to source and residual water treatment plants, but apparently excluding open oceanic waters (9, 16–19); with occasional abundances that exceed those of canonical AOB (9, 20, 21). Nonetheless, it is yet to be shown whether comammox *Nitrospira* truly drives ammonia oxidation in open oligotrophic freshwater and soil environments, their presumed preferred habitat based on genomic and physiological evidence (22, 23).

Here, we sought to identify the active ammonia and nitrite oxidizers in a groundwater-fed RGSF using RNA and DNA stable isotope probing (SIP) coupled to 16S rRNA amplicon sequencing. Lab-scale columns packed with filter material from a full-scale RGSF were fed with effluent water amended with NH_4_^+^ or NO_2_^−^ and with ^13^C labelled or unlabelled HCO_3_^−^ for 15 days in the presence or absence of inhibitors of autotrophic ammonia and nitrite oxidation (24). Our findings indicate that *Nitrospira* drive both ammonia and nitrite oxidation. In addition, several other taxa take up substantial HCO_3_^−^ and their DNA and RNA increases in relative abundance when ammonia was the only energy source provided. This study provides the first *in-situ* evidence of ammonia oxidation by comammox *Nitrospira* in an ecologically relevant complex microbiome.

## Materials and methods

### Sampling sites and procedure

Filter material samples were collected from a rapid gravity sandfilter (biofilter) at Islevbro waterworks (Rødovre, Denmark) in May 2013. The influent and effluent water quality is reported elsewhere (13, 14, 25). Filter material was collected from three random horizontal locations of the biofilter using a hand-pushed core sampler. From the extracted filter material core, the top 10 cm was aseptically segregated on site and stored on ice for further use. A portion was frozen on-site in liquid nitrogen for RNA extraction.

### Column experiments and stable isotope labelling

Experiments were conducted using a continuous-flow lab-scale system consisting of glass columns (2.6 cm diameter, 6 cm long) filled with parent filter material (26.5 cm^3^) as described previously (14). Effluent water from the investigated waterworks was used as the medium in all experiments to avoid interference of other autotrophic processes and approximate full scale conditions.

The experimental design consisted of 4 treatments applied to labeled and unlabeled control columns. The experiments were organized in two phases of 4 columns each; filter material was sampled for DNA and RNA extraction just before the onset of each experimental phase. In the 4 treatments, the influent waters were spiked with: (i) NH_4_^+^ (NH_4_Cl at 1 mg/L N (71 μM); Sigma-Aldrich, 254134), NH_4_^+^ and ATU (N-Allylthiourea at 100 μM, Merck chemicals, 808158), (iii) NO_2_^−^ (NaNO_2_ at 1 mg/L N (71 μM)) Sigma-Aldrich, S2252), (iv) NH_4_^+^ and ClO_3_^−^ (KClO_3_^−^ at 1 mM; 99%, Sigma-Aldrich, 12634) (Table 1) (26).(26). The applied flow rates (40 mL/hr) and influent (NH_4_^+^ or NO_2_^−^) concentrations were set to match the volumetric NH_4_^+^-N loading rates (approx. 1.5 g N/m^3^/hr) experienced by the full-scale parent biofilter (14). Test and control columns were operated for 15 days with continuous feeding to allow sufficient ^13^C label incorporation. Further details are given in *SI Appendix*, SI Materials and Methods.

**Table 1.** Summary of experimental design, bulk ^13^C incorporation, substrate utilization and accumulation levels, and sequenced samples

| Runs | | N source | C source | Inhibitor | $^{13}\text{C}/^{12}\text{C}$ | $\text{NH}_4^+$ <sup>(a)</sup> removal | $\text{NO}_2^-$ <sup>(a)</sup> removal | $\text{NO}_3^-$ <sup>(a)</sup> accretion | Total DNA | Total RNA | SIP |
| --- | --- | --- | --- | --- | --- | --- | --- | --- | --- | --- | --- |
| Run 1<br>Full-scale<br>Inoculum 1 | Col.1 | $\text{NH}_4^+$ | $\text{H}^{13}\text{CO}_3^-$ | - | 279 | 99%±1 | 100%±0 | 88%±32 | + | + | + |
| | Col.2 | $\text{NH}_4^+$ | $\text{HCO}_3^-$ | - | - | 98%±3 | 100%±0 | 82%±28 | + | + | + |
| | Col.3 | $\text{NH}_4^+$ | $\text{HCO}_3^-$ | ATU | - | 19%±15 | 101%±2 | ND <sup>(c)</sup> | + | + | + |
| | Col.4 | $\text{NH}_4^+$ | $\text{H}^{13}\text{CO}_3^-$ | ATU | 54 | 11%±15 | 99%±6 | ND <sup>(c)</sup> | + | + | + |
| Run 2<br>Full-scale<br>Inoculum 2 | Col.5 | $\text{NO}_2^-$ | $\text{H}^{13}\text{CO}_3^-$ | - | 89 | NA <sup>(b)</sup> | 88%±1 | 99%±34 | + | + | + |
| | Col.6 | $\text{NO}_2^-$ | $\text{HCO}_3^-$ | - | - | NA <sup>(b)</sup> | 92%±3 | 62%±36 | + | + | + |
| | Col.7 | $\text{NH}_4^+$ | $\text{HCO}_3^-$ | $\text{ClO}_3^-$ | - | 11%±5 | 70%±44 | ND <sup>(c)</sup> | + | + | + |
| | Col.8 | $\text{NH}_4^+$ | $\text{H}^{13}\text{CO}_3^-$ | $\text{ClO}_3^-$ | 63 | 6%±7 | 97%±28 | ND <sup>(c)</sup> | + | + | + |

<sup>(a)</sup> Removal and accumulation rates were estimated from daily $\text{NH}_4^+$ , $\text{NO}_2^-$ , $\text{NO}_3^-$ measurements
- $\text{NO}_2^-$ removal was calculated based on ammonium removed (except for Col. 5 & 6 where it is based on influent nitrite)
- $\text{NO}_3^-$ accretion was calculated based on ammonium removed (except for Col. 5 & 6 where it is based on nitrite removed)
<sup>(b)</sup> NA: Not Applicable
<sup>(c)</sup> ND: Differences between in and effluent $\text{NO}_3^-$ concentrations were not detectable and accretion could not be calculated

### Analytical methods

Column effluents were sampled daily, filtered (0.2 μm cut-off), frozen and analyzed colorimetrically for NH_4_^+^ and NO_2_^−^ as described in Tatari et al. (26). Colorimetric analysis of ammonium in samples containing ATU underestimated the NH_4_^+^ concentration (26) and thus NH_4_^+^ in these samples was quantified by flow injection analysis (27). NO_3_^−^ was quantified by Ion Chromatography (Dionex, ICS 1500) fitted with a guard column (Dionex, AG 22) and an analytical column (Dionex, ION PAC AS22). NH_4_^+^ removal (%) was calculated by subtracting effluent from influent NH_4_^+^ concentration and normalizing for the influent NH_4_^+^ concentration. NO_2_^−^ removal (%) was calculated as the difference between produced NO_2_^−^ concentration and effluent NO_2_^−^ concentration, after correcting for trace NO_2_^−^ present in the water (ca. 0.3 μM NO_2_^−^) and normalization for the produced NO_2_^−^concentration. The NO_2_^−^ produced by ammonia oxidation was estimated as the difference between influent and effluent NH_4_^+^ concentrations. NO_3_^−^ accumulation (in %) was calculated from the difference between the effluent NO_3_^−^ and influent NO_3_^−^ concentration, normalized for the produced NO_3_^−^ concentration. The NO_3_^−^ produced was estimated as the difference between influent and effluent NH_4_^+^ (or NO_2_^−^ in the case of Col. 5 and 6) concentrations..

### Nucleic acid extraction and stable isotope probing (SIP)

Filter material samples collected from the full-scale biofilter and the sacrificed columns were subject to DNA and RNA extraction. Genomic DNA was extracted from 0.5 g of drained filter material using the MP FastDNA™ SPIN Kit (MP Biomedicals LLC., Solon, USA) according to manufacturer’s instructions. The concentration and purity of extracted DNA was checked by spectrophotometry (NanoDrop Technologies, Wilmington, DE, USA). RNA was extracted from frozen filter material samples (−80 °C) with a MoBio PowerSoil Total RNA Isolation Kit (#12866-25) according to manufacturer’s instructions. The RNA was further purified with a Qiagen AllPrep DNA/RNA Mini Kit (Hilden, Germany) and quantified with a Ribogreen RNA-quantification kit (Invitrogen, Eugene, OR, USA). Extracted rRNA (approximately 650 ng) was mixed well with cesium trifluoroacetate solution to achieve an initial density of 1.790 g/mL before ultracentrifugation at 38,400 rpm for 72 h at 20 °C in Beckmann with a VTi65.2 rotor (28). Centrifuged rRNA gradients were fractionated into 250 μL fractions, the buoyant density of each fraction measured with refractometry and rRNA precipitated from fractions as described previously by Whiteley et al. (29). The concentration of purified RNA was determined using a Ribogreen RNA-quantification kit.

Density gradient ultracentrifugation of DNA isolated from columns and full-scale was performed according to Neufeld et al. (30). Briefly, 1.6 μg of DNA in CsCl with a final density of approximately 1.725 g/mL was subject to ultracentrifugation at 44,800 rpm for 44 h, 20°C in an ultracentrifuge (Beckmann) with a VTi65.2 rotor (Beckmann). Gradients were fractionated into 250 μL fractions, density was determined by refractometry and DNA was recovered by precipitation with PEG. DNA concentration was determined using a Picogreen high sensitivity dsDNA quantification kit (Invitrogen).

### PCR amplification and tag sequencing

RNA purified from density gradient fractions, and from sacrificed column experiments and full scale biofilters were reverse transcribed using reverse primer 1492R with Sensiscript RT kit (Qiagen) according to the manufacturer’s protocol. 10 ng of cDNA or DNA from direct DNA extracts (Table 1) were used to amplify the V3-V4 regions of bacterial 16S rRNA genes using the Phusion (Pfu) DNA polymerase (Finnzymes, Finland) and 16S rRNA gene targeted (rDNA) modified universal primers PRK341F and PRK806R (31). PCR was performed as described in (7). All fractions (a total of 145 fractions; Fig.S1.a) from DNA-SIP and selected fractions (a total of 62 fractions; Supplementary Fig.S1.b) from RNA-SIP experiments were sequenced on an Illumina MiSeq and GS FLX pyrosequencing platform, respectively. Pyrosequencing was applied in a two-region 454 run on a 70-75 GS PicoTiterPlate using a Titanium kit (7); paired-end 16S rRNA amplicon sequencing was done on the Illumina MiSeq platform with MiSeq Reagent Kit v3 (2 × 301 bp; Illumina). All sequencing was performed at the National High-throughput DNA Sequencing Center (Copenhagen, DK).

### Bioinformatic and statistical analysis

All bioinformatic and statistical analyses are described in detail in *SI Appendix*, *SI Materials and Methods*. Briefly, raw 454 sequence data from RNA-SIP samples were quality-checked (denoised) with Ampliconnoise (32) and chimeras were removed with UCHIME (33) using default settings. Raw Miseq Illumina sequence data from DNA-SIP samples were quality-controlled with MOTHUR (34) and chimeras were removed with UCHIME (33) using a reference dataset. Sequence libraries were combined and trimmed to 418 bp. All further sequence analyses were performed in QIIME 1.9.1(35).

A total of six filter steps were applied to identify ammonia and nitrite oxidizing phylotypes (Fig.S2). Detailed steps are described in *SI Appendix, SI Materials and Methods*. OTUs incorporating H^13^CO_3_^−^ were determined by 1. (Filter 1) comparing the mean buoyant density of each OTU in columns with and without ^13^C amendment (36), 2. (Filter 2) identifying OTUs affiliated with genera that are present in both DNA and RNA SIP, and 3. (Filter 3) selecting OTUs with buoyant density shifts higher than genus-specific 90% CIs for buoyant density shifts. The remaining filter steps were applied to assess ammonia and nitrite oxidizing phylotypes. Cross-feeders and taxa performing heterotrophic CO_2_ fixation were largely removed by 4. (Filter 4) excluding OTUs with lower buoyant density shift than the maximum buoyant density shift value of labelled *Nitrosomonas* and *Nitrospira* OTUs, respectively, 5. (Filter 5) selecting the genera which contained OTUs in both RNA and DNA-SIP, and 6. (Filter 6) comparing the labelled OTUs between treatments (NH_4_^+^ fed, NH^+^ - ATU fed, NO_2_^−^ fed, NH_4_^+^-ClO_3_^−^ fed): To identify ammonia oxidizing phylotypes, labelled OTUs in all treatments excluding the one fed only with NH_4_^+^, were removed from the labelled OTU library of NH_4_^+^ fed treatment. To identify nitrite oxidizing phylotypes, labelled OTUs in the treatment fed with ClO3^−^ were removed from the labelled OTU library of the only NO_2_^−^ fed treatment.

As an *additional* step, detected genera were ranked according to the increase in their relative abundance in both total DNA and RNA from the beginning (day 0) to the end (day 15) of the experimental runs.

R codes for all bioinformatics and statistics including the detection of labelled OTUs in DNA and RNA -SIP can be found in https://github.com/ardagulay.

All sequence data have been deposited at NCBI GenBank under Biosample accession numbers from SAMN12227610 to SAMN12227705.

## Results

Four different experimental treatments were designed to identify the microbes involved in ammonia and nitrite oxidation (Table 1): (i) 71 μM of NH_4_^+^ (Cols 1-2), (ii) 71 μM of NH_4_^+^ and 100 μM allylthiourea (ATU) (Cols 3-4), (iii) 71 μM of NO_2_^−^ (Cols 5-6) and (iv) 71 μM of NH_4_^+^and 1 mM NaClO3 (Cols 7-8). All columns were operated with an influent containing 100% ^13^C-labelled or 100% unlabelled bicarbonate for 15 days.

In Col. 1 (H^13^CO_3_^−^) and Col. 2 (HCO_3_^−^), the full-scale conditions were mimicked, with the aim to elucidate the complete *in situ* food web related to nitrification. In Col. 3 (HCO_3_^−^) and Col. 4 (H^13^CO_3_^−^), ATU was used to suppress bacterial ammonia oxidation while feeding at the same NH_4_^+^ loading as in Col. 1 and 2 (26). Complete inhibition of bacterial ammonia oxidation with ATU has been previously observed at ATU concentrations of 8-86 μM (37), while archaeal ammonia oxidation is less sensitive to ATU (38). The mechanism of ATU inhibition in AOB is proposed to be chelation of the Cu^2+^ from the active site in the AMO enzyme (24). To identify taxa associated with nitrite oxidation, NO_2_^−^ was fed to Col. 5 (H^13^CO_3_^−^) and Col. 6 (HCO_3_^−^). In Col. 8 (H^13^CO_3_) and Col. 7 (HCO_3_), ClO_3_^−^ was used to inhibit nitrite oxidation under NH_4_^+^ feeding, with the aim to identify the taxa solely associated with NH_4_^+^ oxidation (26). Chlorate is commonly used as a selective inhibitor for nitrite oxidation, as it is reduced by reverse activity of the nitrite oxidoreductase to the toxic chlorite (ClO_2_^−^)(39, 40).

### Physiological activity

In the 71 μM NH_4_^+^ fed treatments, complete NH_4_^+^ removal (99%) was observed without inhibitor addition, while NH_4_^+^ removal ranged from 11 to 19% with ATU-amendment. (Table 1, Fig.S3.a). Inhibitor addition also significantly reduced overall 13C incorporation. Columns fed with NH_4_^+^, NH_4_^+^-ATU and NO_2_^−^ all had similarly high degrees of NO_2_^−^ removal ranging from 88% to 100%. In the 1 mM ClO_3_^−^ amended columns, NH_4_^+^ removal was severely inhibited (6% to 11%); removal of formed NO_2_^−^ continued (from 70 to 54%), although accumulation of NO_3_^−^ could not be detected. Nitrogen mass balances, based on influent and effluent NH_4_^+^, NO_2_^−^, and NO_3_^−^ concentrations closed for most experimental runs minimizing the possibility of additional nitrogen cycling; N loss was only observed in the ATU supplemented columns (columns 3,4) with ongoing treatment (Table 1; Fig.S3.b-c)

### Detection of ^13^C-labelled taxa from DNA- and RNA- SIP

DNA and RNA, extracted from column samples taken at the end of the experiments, were subject to equilibrium density centrifugation, gradient fractionation and 16S rRNA gene amplification. A total of 147 and 65 gradient fractions from DNA-SIP and RNA-SIP were sequenced using Illumina-Miseq and 454-pyrosequencing platforms, respectively (Fig.S1.a-b). OTUs were defined at 99% similarity, to minimize the effect of microdiversity, as 98.7% and lower similarities represent the taxonomic levels of species, genus and higher (41).

We first examined the incorporation of ^13^C in OTUs in all treatments by comparing replicate columns with H^12^CO_3_^−^ versus H^13^CO_3_^−^ amendment. In DNA-SIP, where all SIP fractions were sequenced, we calculated the average shift in buoyant density of each OTU based on its relative sequence abundance and buoyant density in all fractions (*SI appendix, Eq. 2*). As only selected fractions were sequenced in RNA-SIP, the mean buoyant density of each OTU in treatments with H^12^CO_3_^−^ versus H^13^CO_3_ ^−^ amendment was calculated using the standard deviation of the RNA distribution across the buoyant density gradient (Fig.S4), as described in Zemb *et al*.(36). The buoyant density shift of each OTU was then determined from the calculated mean buoyant density in the replicate columns of each treatment.

Among all detected OTUs (3,364,425), 4,075 and 5,045 in NH_4_^+^ treatment, 4,133 and 5,155 in NH_4_^+^ plus ATU treatment, 4,183 and 706 in NO_2_^−^ treatment, and 44,916 and 52 in the NH_4_^+^ plus ClO_3_^−^ fed treatment showed a buoyant density shift (after Filter 1; Fig.S2)in the DNA- and RNA-SIP experiment, respectively. Only those OTUs that belonged to genera that contained OTUs that were detected as ^13^C-labelled in both RNA and DNA-SIP were retained (Filter 2; Fig.S2). A bootstrap resampling of labelled OTUs within each genus was then used to estimate taxon-specific 90% CIs for the buoyant density shift of a labelled genus (Filter 3; *SI Appendix, M.M* and Fig.S2).

Hence, after the 3^rd^ filter step, 676 (57% DNA, 43% RNA of the total ^13^C-labelled OTUs – NH_4_^+^ fed treatment), 735 (67% DNA, 33% RNA – NH_4_^+^ plus ATU fed treatment), 529 (65% DNA, 35% RNA – NO_2_^−^ treatment) and 43 (83% DNA, 16% RNA – NH_4_^+^ plus ClO_3_^−^ fed treatment) OTUs were retained as significantly labeled. The fractional ^13^C uptake of labelled OTUs was calculated by dividing the DNA and RNA buoyant density shift for each OTU by the total observed buoyant density shift for DNA and RNA, respectively. The abundance of labeled OTUs in the total community were estimated based on total (i.e. non-fractionated) DNA and rRNA extracts collected on day 15 (Fig.S5.a-d).

In the NH_4_^+^-only fed treatment, ^13^C-labelled OTUs affiliated with 17 genera of the *α*-, *β*-, and *γ- Proteobacteria*, *Nitrospira*, *Actinobacteria*, *Latescibacteria*, and *Acidobacteria* (Fig.S5a). Among them, the genus *Nitrospira* had the highest fraction of ^13^C uptake (32% and 1.1% for DNA-SIP and RNA-SIP respectively), and highest relative abundance in the total DNA (26%). *Nitrosomonas* spp. OTUs were also labelled but displayed low levels of ^13^C uptake (0.07% and 0.8% for DNA-SIP and RNA-SIP respectively) and were at low abundance (0.15% and 0.18% in total DNA and RNA, respectively). Labelled ribosomal RNA, an approximation of metabolic activity, was distributed evenly between 5 different ^13^C-labelled genera including *Woodsholea, Blastocella, Subgroup 10 Acidobacteria*, *Pedomicrobium*, and *Sphingomonas* (Fig.S5.a) Although ammonium oxidation was severely inhibited in the NH_4_^+^ plus ATU fed column (Fig.1), OTUs in 15 genera incorporated ^13^C. These were identical to labelled OTUs in the NH_4_^+^ fed treatment with the exception of *OM27*, *Rhizobacter*, *Variovorax* and uncultured representatives of the order *Xanthomonadales*, which were not labelled in the presence of ATU (Fig.S5.b). *Azospira* incorporated H^13^CO_3_^−^ only in the presence of ATU. In the NH_4_^+^ plus ATU fed treatment, *Nitrospira* (10% DNA-SIP, 4.7% RNA-SIP), *Pseudomonas* (3% DNA-SIP, 1% RNA-SIP), *Methyloglobulus* (2.7% DNA-SIP, 2.6% RNA-SIP) and *Blastocatella* (2.6% DNA-SIP, 3.1% RNA-SIP), incorporated the highest fraction of label, while *Sphingomonas* (1.8%) and *Woodsholea* (1.4%) were dominant in the total RNA pool (*Fig.S5.b*). In the NH_4_^+^ plus ClO_3_^−^ amended columns, where both ammonium and nitrite oxidation were suppressed, *OM27* (2.7% DNA, 2.6% RNA) and *Woodsholea* were the only taxa that assimilated significant amounts of HCO_3_^−^ (Fig.S5.c).

**Fig. 1.**
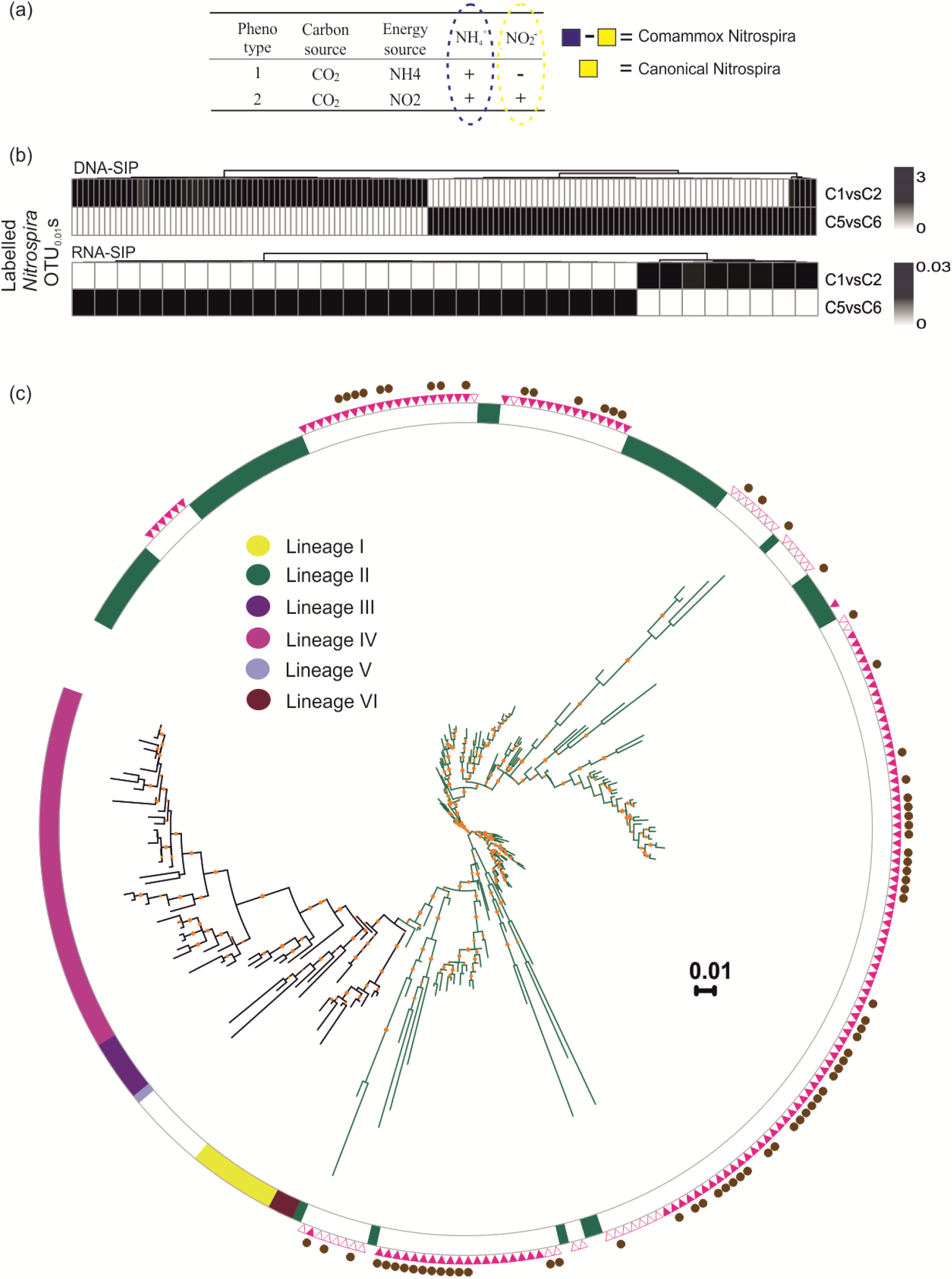
(a) Approach to identify ammonia and nitrite oxidizing *Nitrospira* (b) Heatmap of all identified labelled *Nitrospira* OTUs in columns fed with NH_4_^+^ and NO_2_^−^, (c) 16S rRNA based phylogenetic tree of all identified labelled *Nitrospira* OTUs and published *Nitrospira* strains with known lineages. Open and filled triangles represent *Nitrospira* OTUs identified by RNA and DNA-SIP, respectively. Comammox *Nitrospira* sequences are indicated by filled circles. Bootstrap values >60% are shown by orange dots. The scale bar represents 0.01 substitutions per nucleotide position.

In the NO_2_^−^ fed columns, 12 genera, belonging to *α-Proteobacteria* (10% of H^13^CO_3_^−^uptake in DNA-SIP, 23% of H^13^CO_3_^−^uptake in RNA-SIP), δ-Proteobacteria (1% DNA-SIP, 21% RNA-SIP) and *γ-Proteobacteria* (6% DNA-SIP, 11% RNA-SIP), *Nitrospira* (71% DNA-SIP, 15% RNA-SIP), *Actinobacteria* (3% DNA-SIP, 7% RNA-SIP), *Latescibacteria* (0.5% DNA-SIP, 5% RNA-SIP), and *Acidobacteria* (6% DNA-SIP, 11% RNA-SIP), were labelled (after Filter 3; *Fig.S5.d*). *Nitrospira* had the highest number of labelled OTUs (96) and was responsible for the majority of the H^13^CO_3_^−^ uptake. After application of Filters 4 through 6, only *Nitrospira* OTUs were retained and hence identified as the sole nitrite oxiders.

### Hypothesis: Nitrospira is an active ammonium oxidizer

H^13^CO_3_^−^ was incorporated by *Nitrospira* in treatments fed with NH_4_^+^, NH_4_^+^ plus ATU and NO_2_^−^ (after Filter 3, Fig.S5.a-d). The observed labelling in individual treatments does not indicate whether labelled *Nitrospira* OTUs are capable of ammonia oxidation because both ammonium and nitrite oxidation occur in NH_4_^+^ treatment. We therefore performed a binary comparision between labelled *Nitrospira* OTUs detected in the NH_4_^+^ versus NO_2_^−^ fed treatments (Fig.1a). We assume that the labelled *Nitrospira* OTUs in NH_4_^+^ fed treatment would include both comammox and nitrite oxidizing *Nitrospira*, while the NO_2_^−^ fed treatment would exclude comammox *Nitrospira* based on observation that comammox *Nitrospira* growth is not supported by oxidation of environmental nitrite in the absence of ammonia(11).

Heatmaps of labelled *Nitrospira* OTUs (Fig.1.b) reveal that 8 (24%) and 70 (51%) OTUs are uniquely labelled in NH_4_^+^ fed treatment at the RNA and DNA levels, respectively, indicating that several ammonium oxidizing *Nitrospira* strains actively assimilating H^13^CO_3_^−^. A high number of labelled *Nitrospira* OTUs are unique to the NH_4_^+^ fed column, and few *Nitrospira* OTUs are shared between the NH_4_^+^ and NO_2_^−^ fed columns suggesting that most comammox *Nitrospira* do not readily switch from ammonium oxidation to solely nitrite oxidation. A large number of OTUs were also uniquely labelled in the NO_2_^−^ amended columns (25 in RNA and 66 in DNA),which suggests that, in the NH_4_^+^ fed treatment, the produced NO_2_^−^ was not sufficient to achieve labelling of nitrite-oxidizing *Nitrospira* due to complete nitrification by commamox *Nitrospira*.

Based on their ^13^C labelling in NH_4_^+^ and NO_2_^−^ fed treatments, 78 (8 in RNA-SIP and 70 in DNA-SIP) and 96 (25 in RNA-SIP and 71 in DNA-SIP) *Nitrospira* OTUs were identified as complete ammonia- and nitrite-oxidizing, respectively (Fig.1c). All labelled *Nitrospira* belonged to lineage II, which comprises both comammox and non-comammox types. No clear branching between comammox and nitrite oxidizing phylotypes was observed from the tree topology.

### High H^13^CO_3_^−^ incorporation and growth by other bacteria

In our previous 16S rRNA amplicon-based analysis of the same and related RGSF communities, members of the *Rhizobiales* (*α*-*proteobacteria)*, and *Acidobacteria* were consistently more abundant than *Nitrosomonas* where NH_4_^+^ is thought to be the largest source of energy available for microbial growth (7). We observed that some of these taxa incorporated ^13^HCO_3_^−^ both in RNA- and DNA-SIP (after Filter 6; Fig. 2a). In addition, several OTUs displayed higher buoyant density shifts than *Nitrosomonas* in NH_4_^+^-fed columns (Fig.S5.a). Finally, these OTUs also increased in relative abundance in both DNA and RNA over the course of the experiment (Fig.3c).

**Fig. 2.**
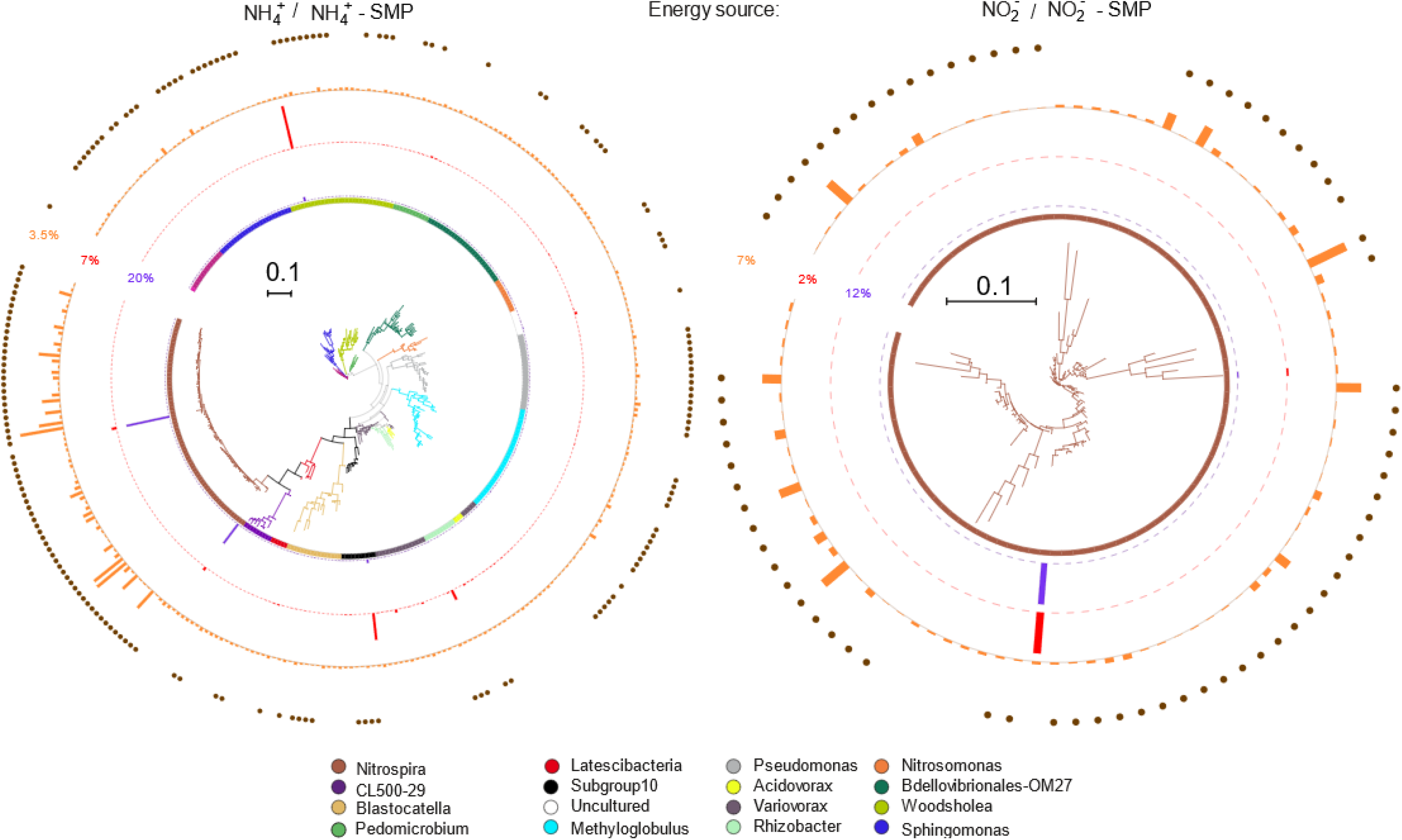
16S rRNA based phylogenetic tree showing phylotypes incorporating H^13^CO_3_^−^ in DNA-SIP and RNA-SIP experiments selective for putatively (a) ammonium and (b) nitrite oxidation. Peak heights on circles represent (i) relative abundance in total DNA (purple) and (ii) total RNA (red) after 15 days, and ^13^C label percentage (orange). The outer ring represents the OTUs retrieved from DNA-SIP (filled circles) or RNA-SIP (no circles). The scale bar represents 0.10 substitutions per nucleotide position.

Only *Nitrospira* was associated with *both* ammonium and nitrite oxidation (after Filter 6; Fig.2a-b). Among ^13^HCO_3_^−^ incorporating taxa in the presence of NH_4_^+^, Subgroup 10 (*Acidobacteria*), *Nitrospira (Nitrospira)*, *Pedomicrobium*, (α-*Proteobacteria*), *Rhizobacter* and *Acidovorax* (*β*-*Proteobacteria*) displayed higher shifts in relative RNA and DNA abundance compared to *Nitrosomonas* (Fig 3c). With the exception of *Pseudomonas* (Fig.S6 *A*), ^13^C-labelled genera in SIP columns (Fig.S5.a-d) differed from the dominant genera in the feedwater, itself the effluent from the full-scale biofilter. Hence, invasion from feed water communities did not cause the increased relative abundance of *Subgroup 10 Acidobacteria*, *Pedomicrobium*, *Rhizobacter*, and *Acidovorax*.

**Fig. 3.**
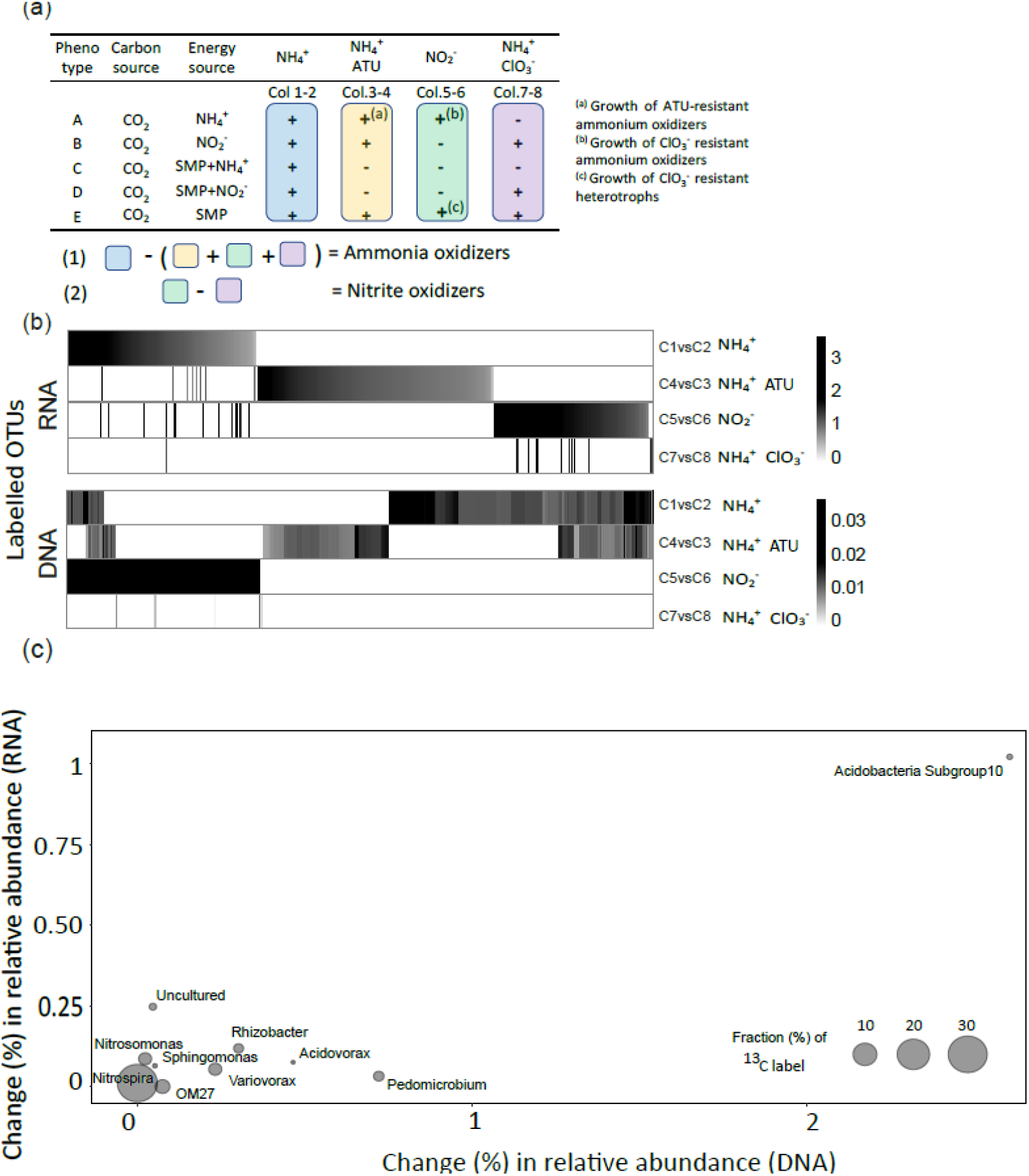
(a) Approach (Filter 6; SI Methods, Fig. S3) to identify putative ammonia and nitrite oxidizing phylotypes: (b) Heatmap of OTUs significantly labelled under NH_4_^+^, NO_2_^−^, NH_4_^+^ plus ATU, and NH_4_^+^ plus ClO_3_^−^ treatment (c) Fold-change in relative abundance in community DNA and RNA for taxa identified as putative ammonia oxidizers (as shown also on Fig 2A).

A metagenome, obtained from the same parent biofilter within the same year (2013, (Palomo et al. 2016)) was further examined for its content of genes that are not canonical AOB *amoA*, canonical AOA *amoA*, *or* comammox *Nitrospira amoA*, nor methanotrophic *pmoA* (Palomo et al. 2016; Fig.S7) but showed homology with genes encoding AMOA protein family fragments from putative heterotrophic nitrifiers (PF05145, IPR017516 Pfam and InterPro database, respectively). Twenty-nine unique *amoA* gene fragments matching the PF05145 model were aligned with reference putative heterotrophic *amoA* gene fragments (Fig. S7). However, no *amoB* and *amoC* genes were found on any of the contigs carrying these atypical putative *amoA* gene fragments (Fig.S8); furthermore no similar gene synteny was detected between the other genes of PF05145 *amoA* containing contigs and our metagenome *amoA* containing contigs (Fig.S8).

The genus *Nitrospira* was the only taxon associated with nitrite oxidation (Fig. 2.b).

## Discussion

Stable isotope probing has previously been used to identify active nitrifiers in sediments (42–44) and soils (45–48). In most studies, either DNA-SIP (45) or RNA-SIP (49) are applied individually; yet both are important to identify key catalysts(48). While DNA-SIP detects isotope incorporation into dividing cells, RNA-SIP detects active potentially slow- or non-growing cells (50). By coupling SIP with next generation sequencing (NGS) we improved taxonomic resolution and differentiated phylotypes in taxa with high microdiversity.

Here, we examined the assimilation of H^13^CO_3_^−^ coupled to nitrification in a RGSF using both RNA- and DNA-SIP. OTUs incorporating ^13^C isotope label in the different treatments were unambiguously identified as those displaying significant buoyant density shifts between the H^12^CO_3_^−^ and the H^13^CO_3_^−^ replicates, and were detected at high phylogenetic resolution (>99% pairwise identity (41)). A total of 200 gradient fractions were processed with sample size equalization.

Our results provide the first *in-situ* physiological evidence of ecologically relevant NH_4_^+^ oxidation by comammox *Nitrospira* in any environment. Other reports of i*n situ* activity are inferred from bulk observations (ammonium removal when comammox *Nitrospira* are more abundant than AOBs or AOAs (20, 51)) or from comammox-specific *amoA* transcript analysis (52). Our former observations that *Nitrospira* was more abundant than *Nitrosomonas* and the discovery that *Nitrospira* harbours the complete nitrification pathway in a full-scale RGSF microbiome (8, 10, 11) is thus directly linked to the ammonia oxidizing activity of *Nitrospira* in this environment. *Nitrospira* was the only genus incorporating H^13^CO_3_^−^ in both NH_4_^+^ fed and NO_2_^−^ fed treatments, indicating that *Nitrospira* is the only genus oxidizing both environmental NH_4_^+^ and NO_2_^−^ in this system.

Ammonia and nitrite oxidizing phylotypes of *Nitrospira* were compared phylogenetically; the resulting 16S rRNA tree topology shows no clear evolutionary separation of comammox and canonical *Nitrospira*. This is in line with previous studies that show that comammox *Nitrospira* are not evolutionarily distant from known canonical *Nitrospira* (>99% 16S rRNA nucleotide identity) (10). Furthermore, our phylogenetic analysis shows that the labelled – both ammonia oxidizing and nitrite oxidizing - *Nitrospira* OTUs, branch within *Nitrospira* sublineage II, as reported in previous studies (8, 10, 11, 53). We did not identify the *amoA* clade affiliation of the active comammox *Nitrospira* phylotypes in this study, although separate investigations on this and related RGSF have indicated a predominance of *amoA* clade B comammox *Nitrospira* (9, 23).

It remains unclear whether comammox *Nitrospira* can switch between modes of ammonia and nitrite oxidation. However, the large numbers of *Nitrospira* phylotypes which were exclusively labelled in the NH_4_^+^ versus NO_2_^−^ fed columns, respectively, suggests that comammox *Nitrospira* may not prefer to oxidize external NO_2_^−^ alone, in agreement with observations in *Can*. N. inopinata (10, 11).

Although ClO_3_^−^ is a well-known competitive inhibitor of nitrite oxidoreductase (54), strong inhibition of *both* ammonium and nitrite oxidation was observed in NH_4_^+^ plus ClO_3_^−^ fed columns. We expect that the inhibition of nitrite oxidation in comammox *Nitrospira* would be caused by ClO_3_^−^ reduction to the toxic ClO_2_^−^, which would negatively affect overall metabolism in these organisms including ammonia oxidation. Hence, it appears that the inhibitory effect of ClO_3_^−^ on ammonium oxidation provides preliminary support for the contribution of comammox *Nitrospira* to ammonia oxidation, as we observed before (26). In addition, ClO_2_^−^ may contribute to inhibition of other ammonia oxidizers as observed before (55). PTIO has also been documented as a potent inhibitor of ammonia oxidation in *N. inopinata*, although its selectivity is unclear (56). ATU significantly suppressed ammonia oxidation in NH_4_^+^-ATU fed treatments, although the taxa assimilating ^13^HCO_3_^−^ did not change significantly compared to the NH_4_^+^ fed treatment, excluding *Azospira*. This similarity in labelled taxa in the ATU-fed treatment may indicate that the AMO of the major ammonia oxidizers in this environment may be less sensitive to ATU than AOB at the given concentrations, as previously observed for AOA (57).

No archaeal taxa were ^13^C-labelled in any of the columns although archaeal ammonium oxidizers (AOA) are present, be it at much lower abundance than *Nitrospira* (100 to 1000 fold), in the RGSF used in this study (8, 13, 58). Columns were fed 71 μM NH_4_^+^ to mimic full-scale conditions (14), while bottom layers of the full-scale biofilter receive very low ammonium concentrations due to removal in the top layers (14, 15). The absence of AOA in the ^13^C-labelled taxa may be due to their low initial abundances, or the elevated concentrations applied during the experiment, as AOA may thrive better in conditions of reduced energy supply consistent with their elevated abundance at bottom layers of the examined RGSF (59–62); even though a recently isolated comammox strain *N. inopinata* displayed higher NH_4_^+^ affinity than many of the characterized AOAs (22).

After strong filters were applied to remove heterotrophic OTUs, we retain the following taxa with substantial H^13^CO_3_^−^ incorporation in the NH_4_^+^ fed treatments: *Subgroup10, Pedomicrobium, Rhizobacter*, and *Acidovorax*; these taxa also show a greater fold-change in relative abundance in DNA and RNA during the experiment than *Nitrosomonas* and *Nitrospira* OTUs (Fig.3c; Fig.S9). Most heterotrophic microbes can engage in CO_2_ fixation via carboxylation reactions (63, 64). However, CO_2_ fixation via anaplerotic metabolism (65) typically results in only 3-8% of the cellular carbon assimilated by heterotrophs, which would be insufficient for label detection by DNA-SIP (63, 66). Thus, heterotrophic carbon fixation would not explain the higher H^13^CO_3_^−^ incorporation extents (higher density shifts, See Fig.2, TableS1, S2) relative to *Nitrosomonas* OTUs, known ammonium oxidizers. Furthermore, the cellular mass and activity supported by cross-feeding decay products from autotrophs (67) would be significantly less than the chemolithoautotrophic biomass and activity itself. Thus, the observation of ^13^C-labelled genera with higher buoyant density shifts and higher fold-changes in DNA and RNA abundance shifts (Fig.2, Fig.3) compared to *Nitrosomonas* and *Nitrospira*, are difficult to explain by cross-feeding alone.

Can a plausible explanation for the high H^13^CO_3_^−^ assimilation of these taxa be nitrification? An earlier metagenome from the same parent material revealed the presence of putative *amoA* genes that could not be classified as *amoA* from canonical AOB, canonical AOA, or comammox *Nitrospira* ((8); Fig.S7); yet were phylogenetically related to the PF05145, purported to contain AMOA encoding genes in heterotrophic bacteria (68). In addition, the phylogeny of 10/30 of these aberrant *amoA* genes indicated their presence, among others, in *Hyphomicrobiaceae* and *Comamonadaceae*. Of the highly labeled taxa (in both RNA-SIP, and DNA-SIP) in the NH_4_^+^ fed treatment, *Pedomicrobium*, and *Acidovorax* (but not *Subgroup10* or *Rhizobacter*) belong to the *Hyphomicrobioaceae* and *Comamondaceae*. While it is tempting to speculate that we have identified novel ammonia oxidizing bacteria, we are unable to identify additional *amo* genes that would constitute a complete *amo* operon on any of the metagenomic contigs. In addition, recent doubt has been cast on the assignment of PF05145 as encoding a putative ammonia monooxygenase (69), and careful physiological or genomic evidence of heterotrophic nitrification remains elusive (70). On the other hand, some of the Acidobacterial MAGs that were retrieved from the studied RGSF metagenome, contained CO_2_ fixation pathways (8).

The second step of nitrification, the oxidation of nitrite to nitrate, is known to be performed by nitrite-oxidizing chemolithoautotrophs such as *Nitrotoga*, *Nitrospina*, *Nitrobacter*, *Nitrolancea* and *Nitrospira* (71–73), which use nitrite oxidoreductase (NXR) as the key enzyme. The known autotrophic nitrite oxidizer *Nitrospira* was identified as the only active nitrite oxidizer in the studied system.

In summary, comammox *Nitrospira* and *Nitrosomonas* are the chemolithoautotrophic drivers of ammonia oxidation in the groundwater fed biofilter and comammox *Nitrospira* make the greatest contribution to ammonia oxidation. This finding raises the question of whether comammox *Nitrospira* is an equally important ammonia oxidizer in other environments. AOA did not significantly contribute to nitrification and *Nitrospira* were the only nitrite oxidizers identified in this environment. Hence, we provide the first *in-situ* evidence of ecologically relevant ammonia oxidation by comammox *Nitrospira* in a complex microbiome and document an unexpectedly high H^13^CO_3_^−^ uptake and growth of *Proteobacterial* and *Acidobacterial* taxa under ammonium selectivity.

## Acknowledgments

We thank Dr. Hannah Sophia Weber and Heidi Grøn Jensen for support with SIP, and Alex Palomo for feedback on the manuscript. This research was supported by The Danish Council for Strategic Research (Project DW Biofilter), the European Commission (MERMAID-ITN; An initial training network funded by the PeopleProgramme - Marie Skodowska-Curie Actions-of the European Union’s Seventh Framework Programme FP7/2007-2013/ under REA grant agreement n1607492) and the VILLUM foundation (Project Expa-N, 13391).

