## Supplementary Figures and Materials for "DNA and RNA-SIP reveal *Nitrospira spp.* as key drivers of nitrification in groundwater-fed biofilters"

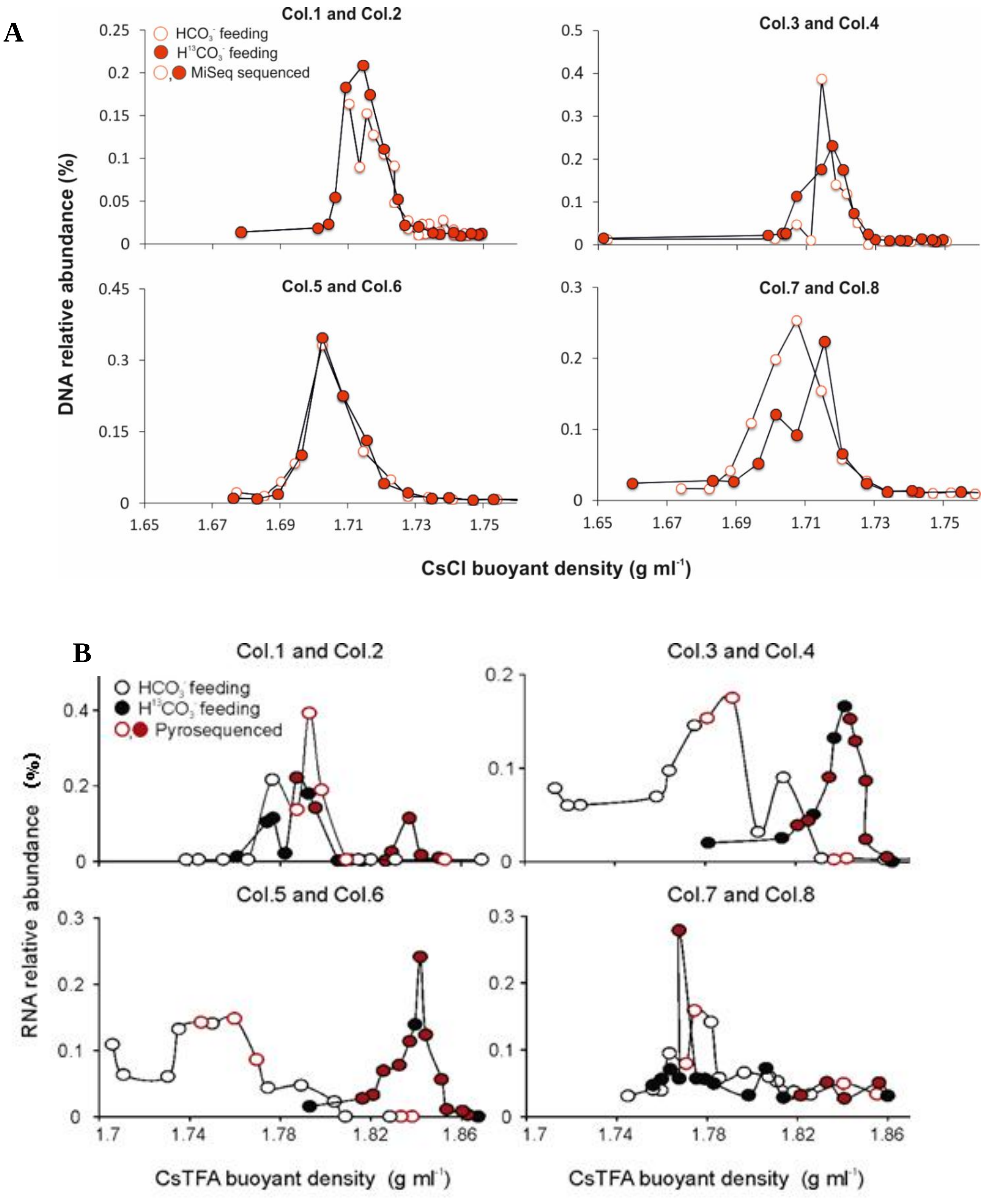


**Fig.S1** (A) Profile of DNA concentration across the CsCl buoyant density gradient obtained from DNA-SIP fractions after 15 days of operation (B) Profile of RNA concentration across the CsTFA buoyant density gradient obtained from RNA-SIP fractions after 15 days of operation

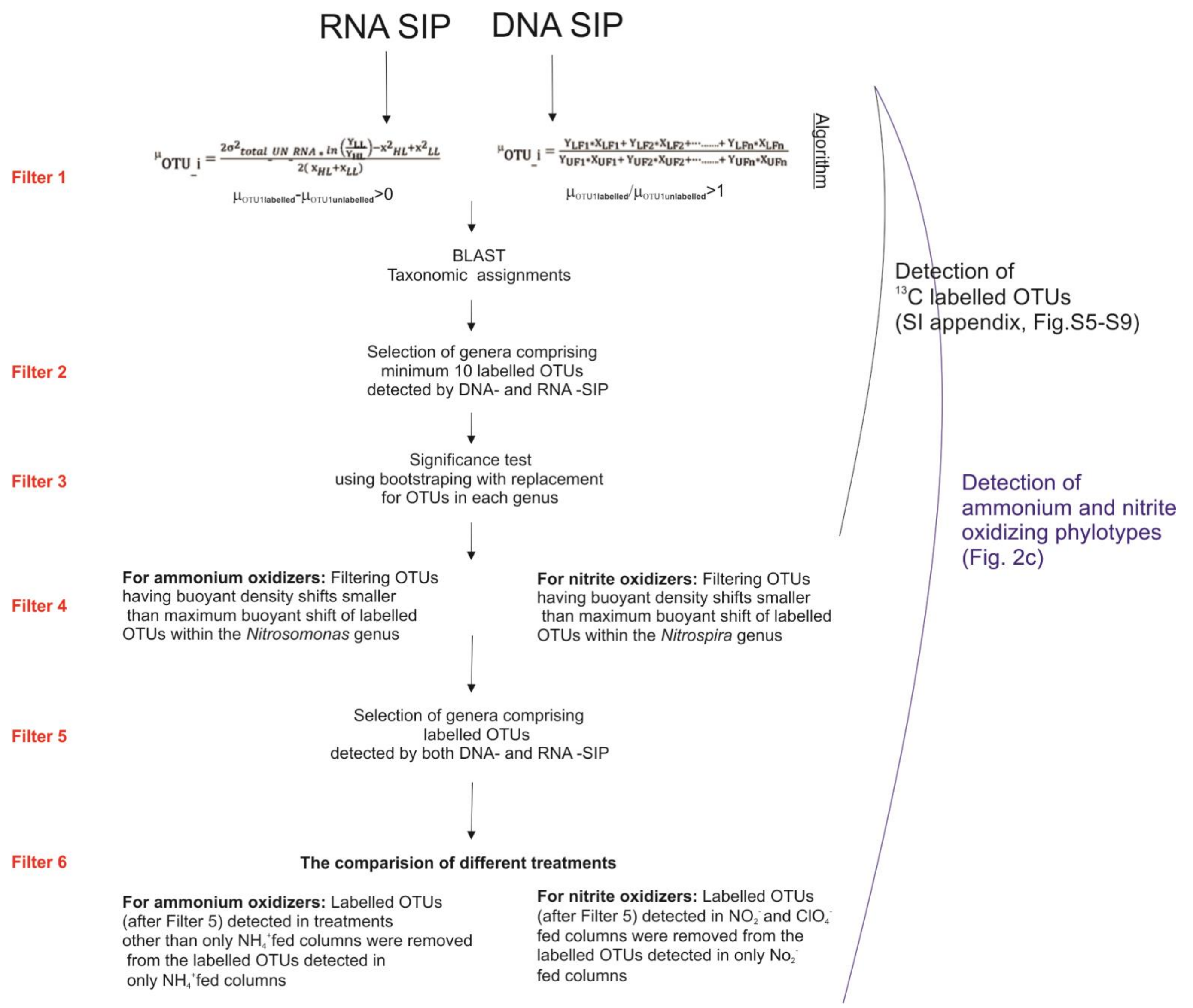


**Fig.S2** Sequential work-flow to detect putative ammonium and nitrite oxidizers using RNA and DNA SIP. Detailed explanation of each filter step is given in Supplementary Materials and Methods.

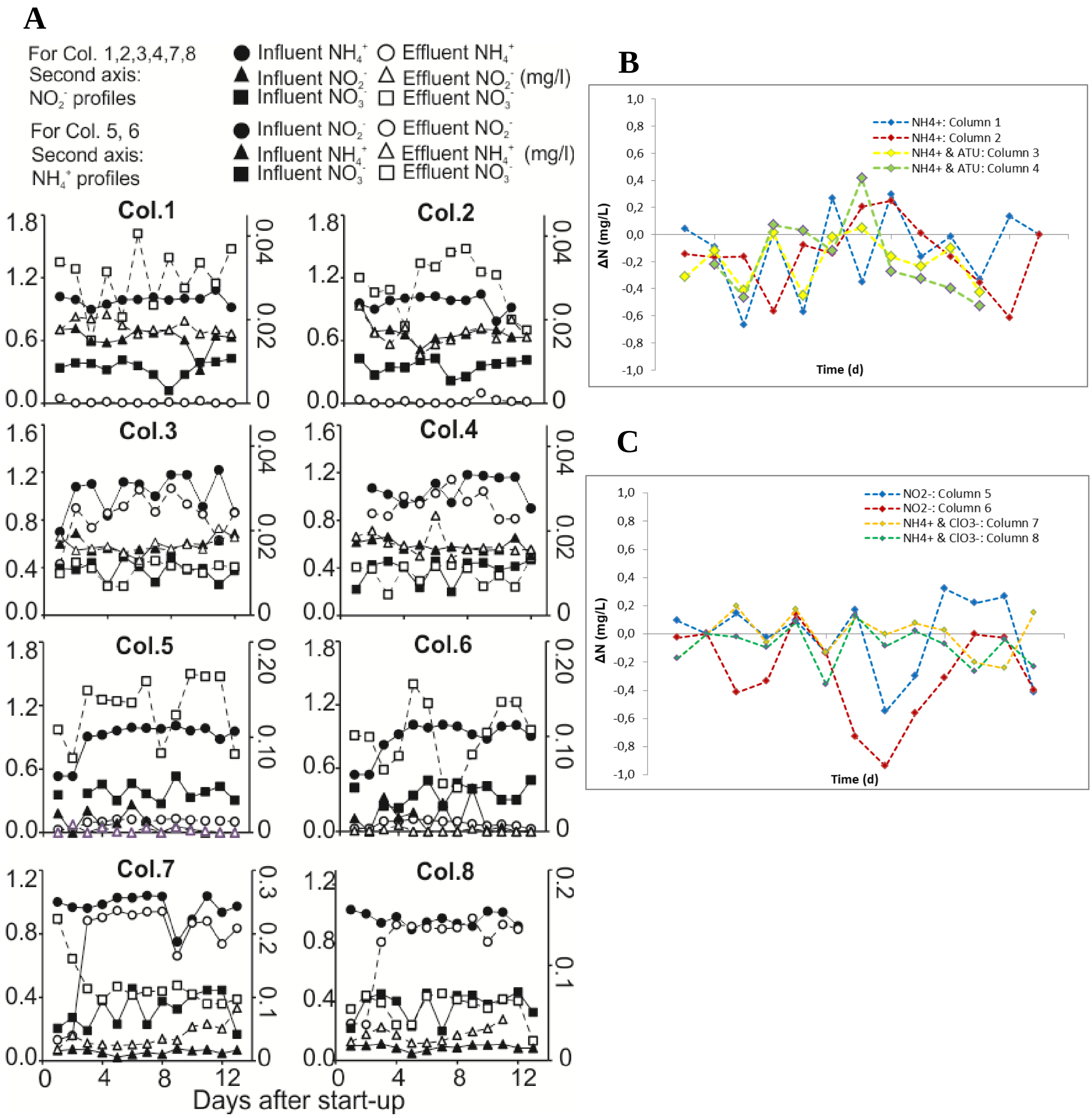


**Fig.S3** (A) Changes in $NH_4^+$, $NO_2^-$, and $NO_3^-$ concentrations in all columns, (B-C) Difference in total N concentration in the influent and the effluent (σ= effluent N-influent N) calculated with the equation 1 given in the *SI* M.M. (B) σ values as a function of operation days for column 1,2,3,4 ($NH_4^+$ and $NH_4^+$ plus ATU treatment), (C) σ values as a function of operation days for column 5,6,7,8 ($NO_2^-$ and $NH_4^+$ plus $ClO_3^-$ treatment). The significance of the difference between influent and effluent total N was evaluated using a 2-tailed t-test (significance level 0.05). Differences were rejected for columns 1-2 and 5-8. In columns 3 and 4, the mass balance indicated N loss during the last sampling points.

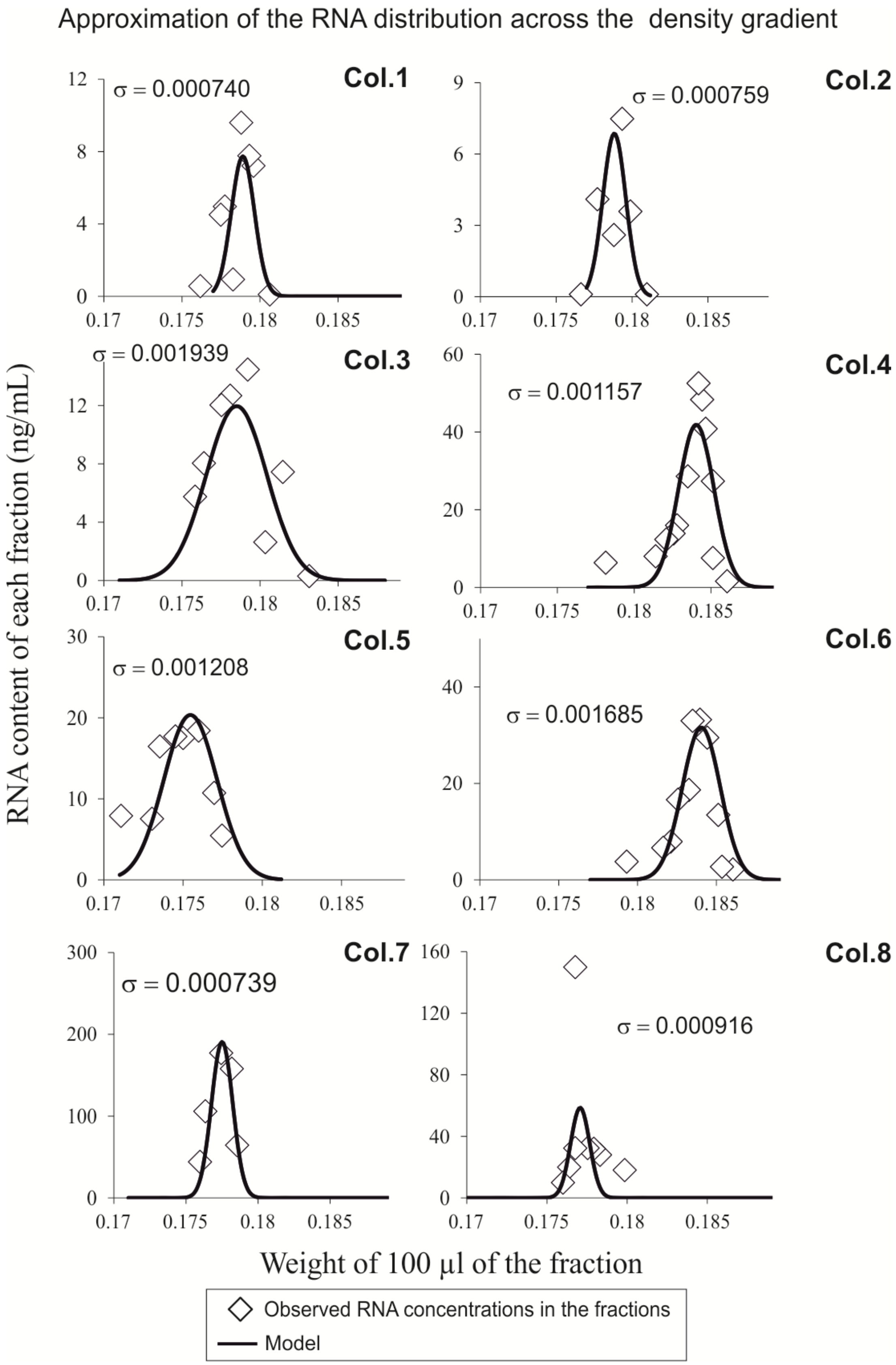


**Fig.S4** Simulated and actual RNA concentration profiles across fractions based on a normal distribution, following the procedure by Zemb *et al.* (1)

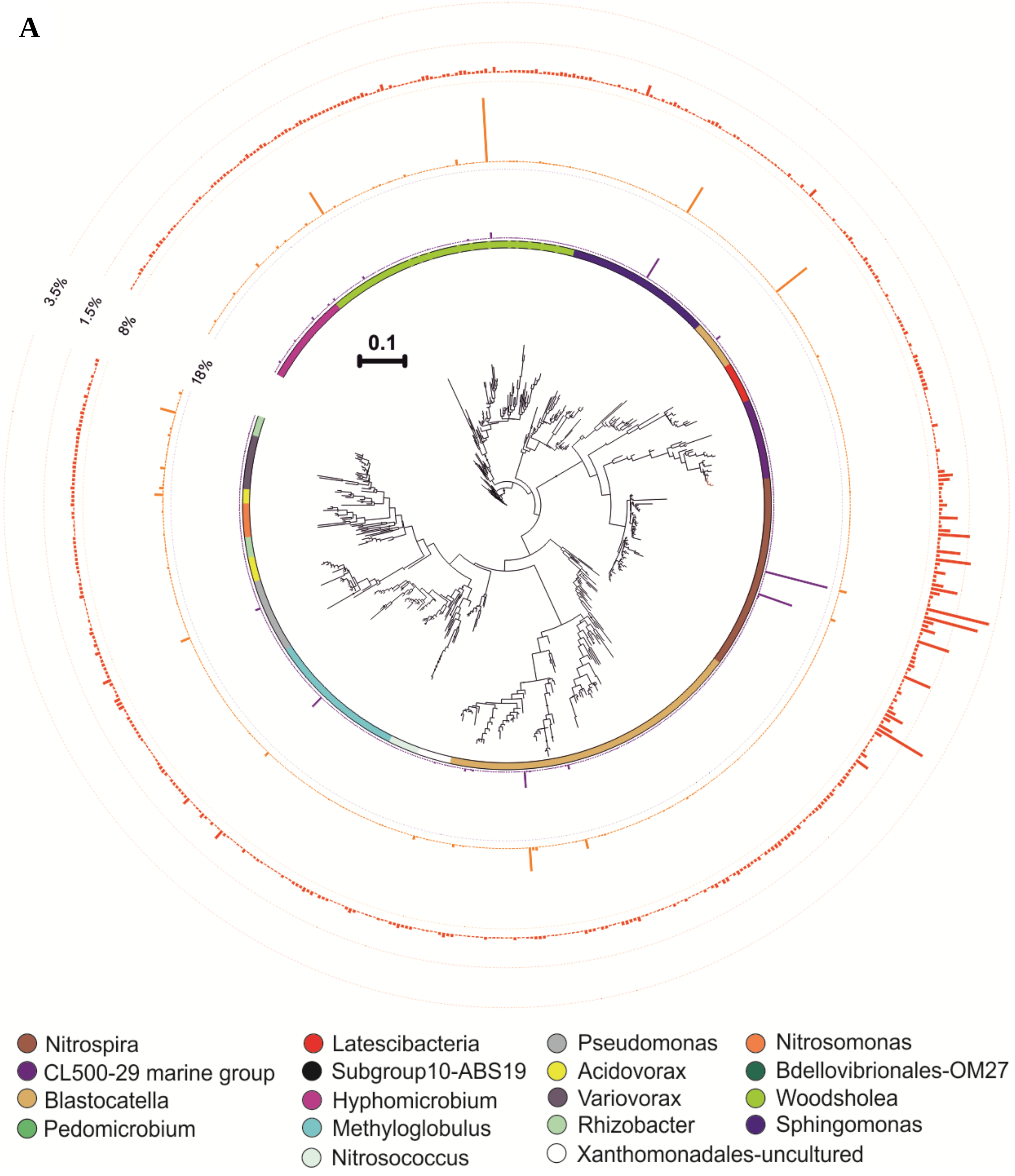
A
0.1
3.5%
1.5%
8%
18%
Nitrospira
CL500-29 marine group
Blastocatella
Pedomicrobium
Latescibacteria
Subgroup10-ABS19
Hyphomicrobium
Methyloglobulus
Nitrosococcus
Pseudomonas
Acidovorax
Variovorax
Rhizobacter
Xanthomonadales-uncultured
Nitrosomonas
Bdellovibrionales-OM27
Woodsholea
Sphingomonas

**B**

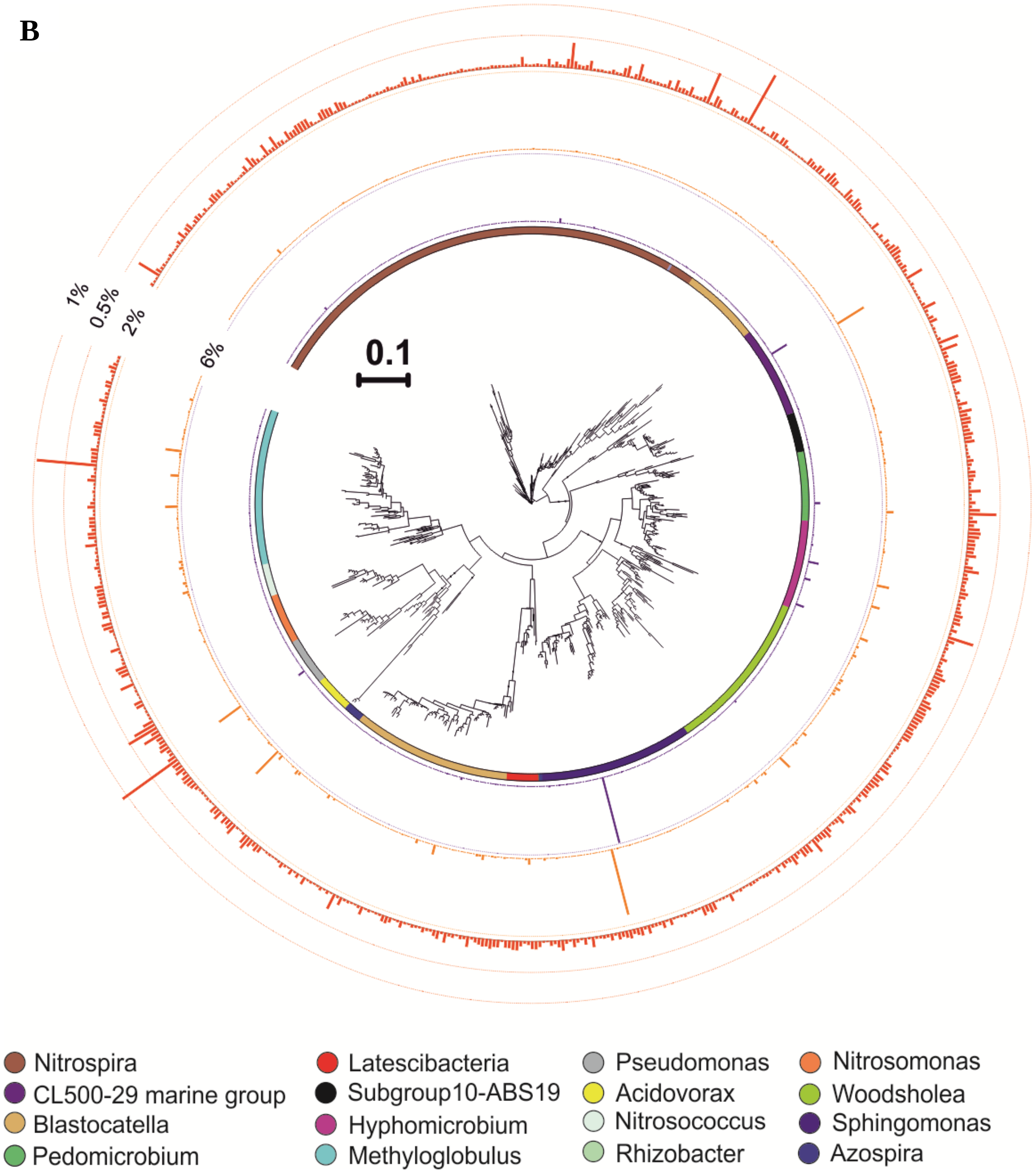
1%
0.5%
2%
6%
0.1
Nitrospira
CL500-29 marine group
Blastocatella
Pedomicrobium
Latescibacteria
Subgroup10-ABS19
Hyphomicrobium
Methyloglobulus
Pseudomonas
Acidovorax
Nitrosococcus
Rhizobacter
Nitrosomonas
Woodsholea
Sphingomonas
Azospira

C

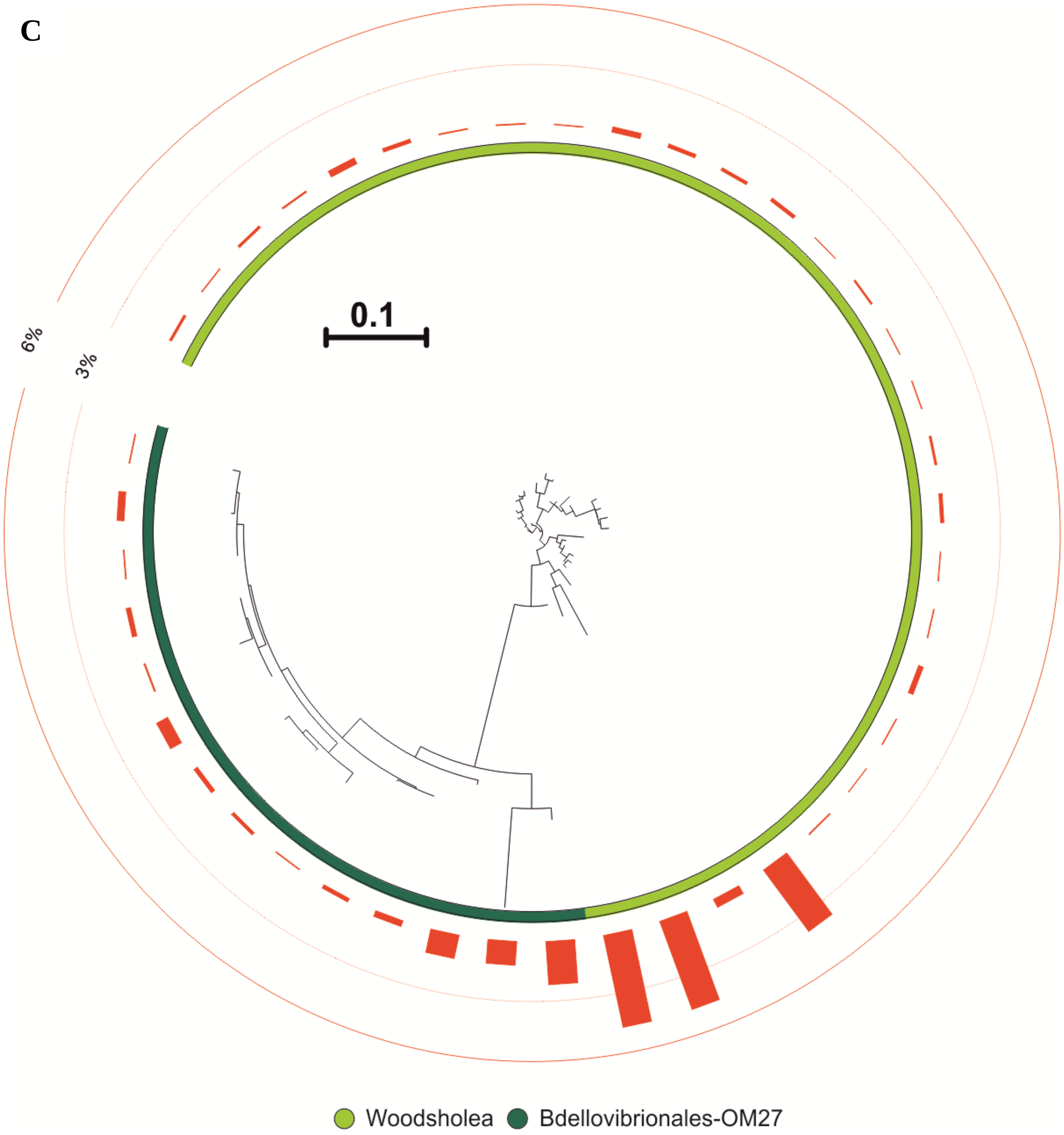

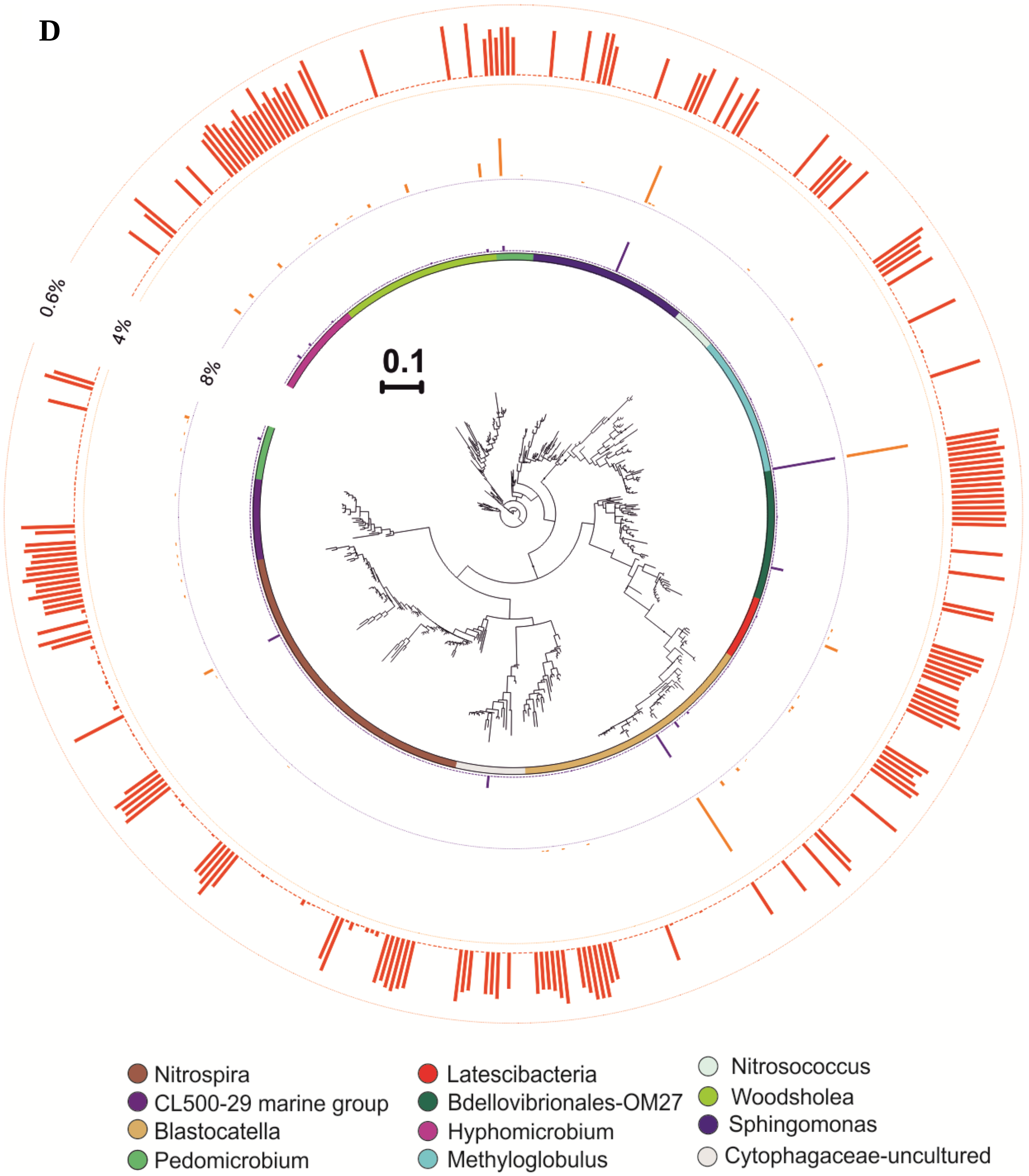


**Fig.S5 .** 16S rRNA based phylogenetic tree of OTUs incorporating $HCO_3^-$ in DNA and RNA-SIP experiments of treatements consisting of (A) solely $NH_4^+$, (B) $NH_4^+$ plus ATU, (C) $NH_4^+$ plus $ClO_3^-$ (D) solely $NO_2^-$. Peak heights on circles represent (i) relative abundance in total DNA (purple) and (ii) total RNA (red) after 15 days, and (iii) $^{13}C$ label percentage (orange). The scale bar represents 0.10 substitutions per nucleotide position.

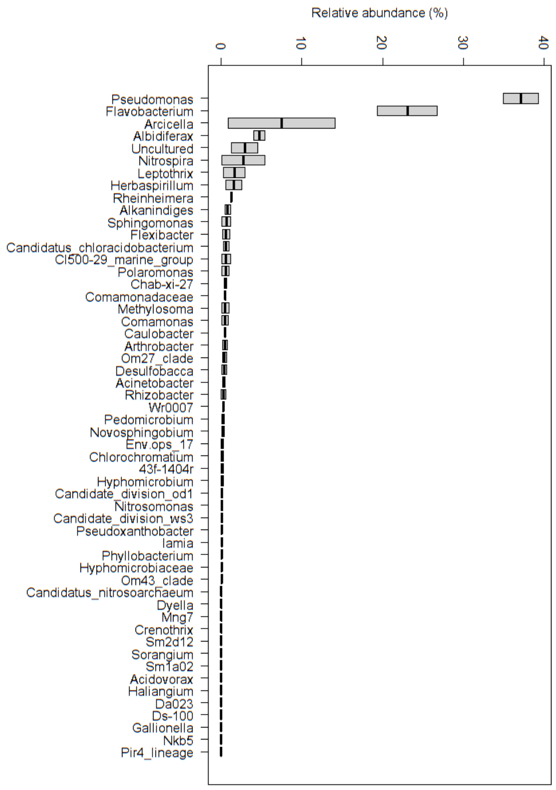


**Fig.S6 A** 16S rDNA based relative abundances of dominant genera (in %) in the communities in the feedwater to the test columns.

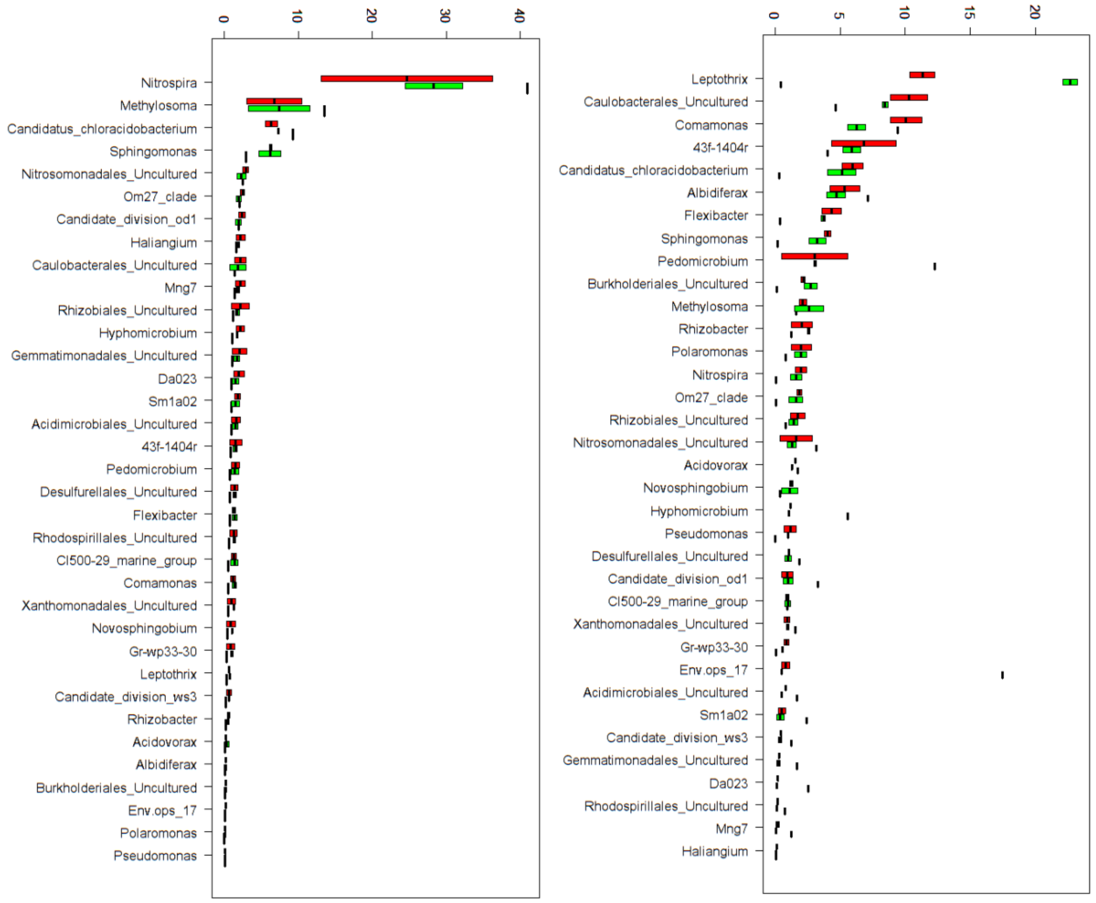


**Fig. S6 B** 16S rDNA (left) and 16S rRNA (right) based relative abundance of dominant genera (in %) in communities extracted from the sand filter material at the onset (black lines) and after 15 days in columns 1 and 2 (green boxes) and columns 3 and 4 (red boxes).

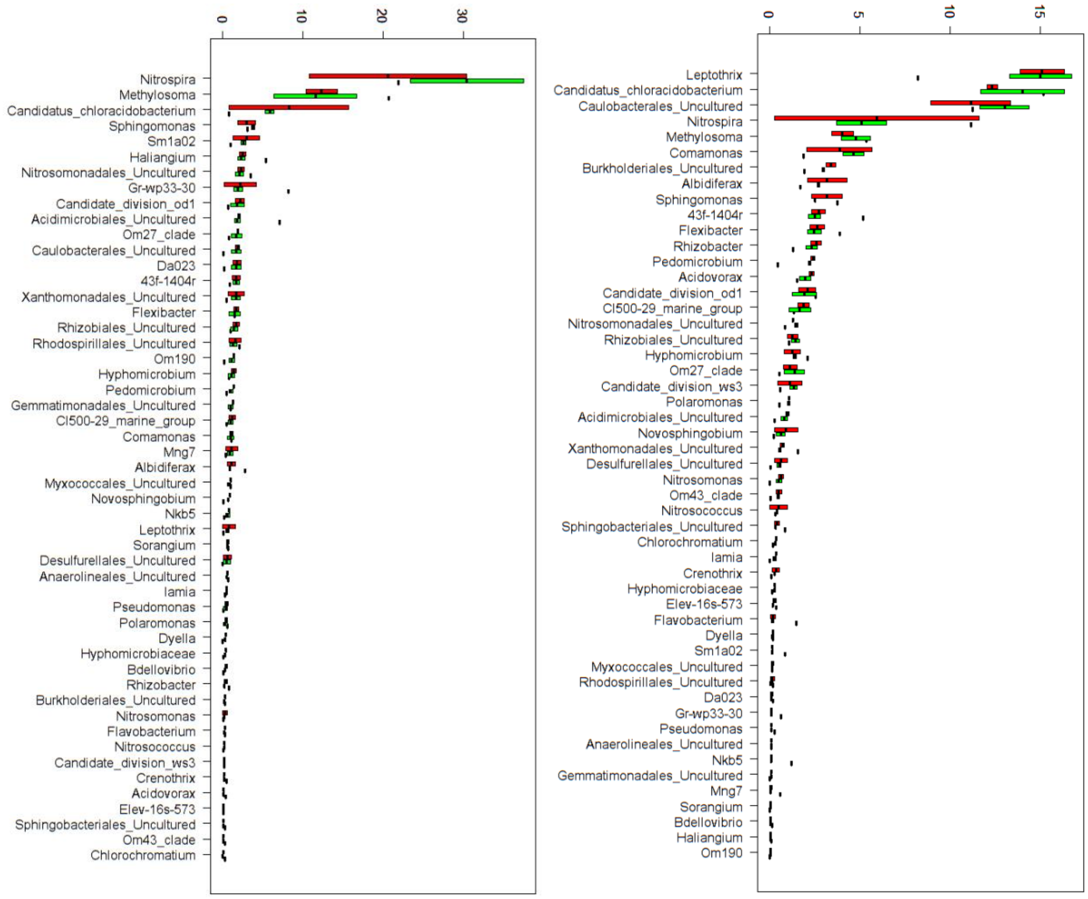


**Fig. S6 C** 16S rDNA (left) and 16S rRNA (right) based relative abundance of dominant genera (in %) in communities extracted from the sand filter material at the onset (black lines) and after 15 days in columns 5 and 6 (green boxes) and columns 7 and 8 (red boxes).

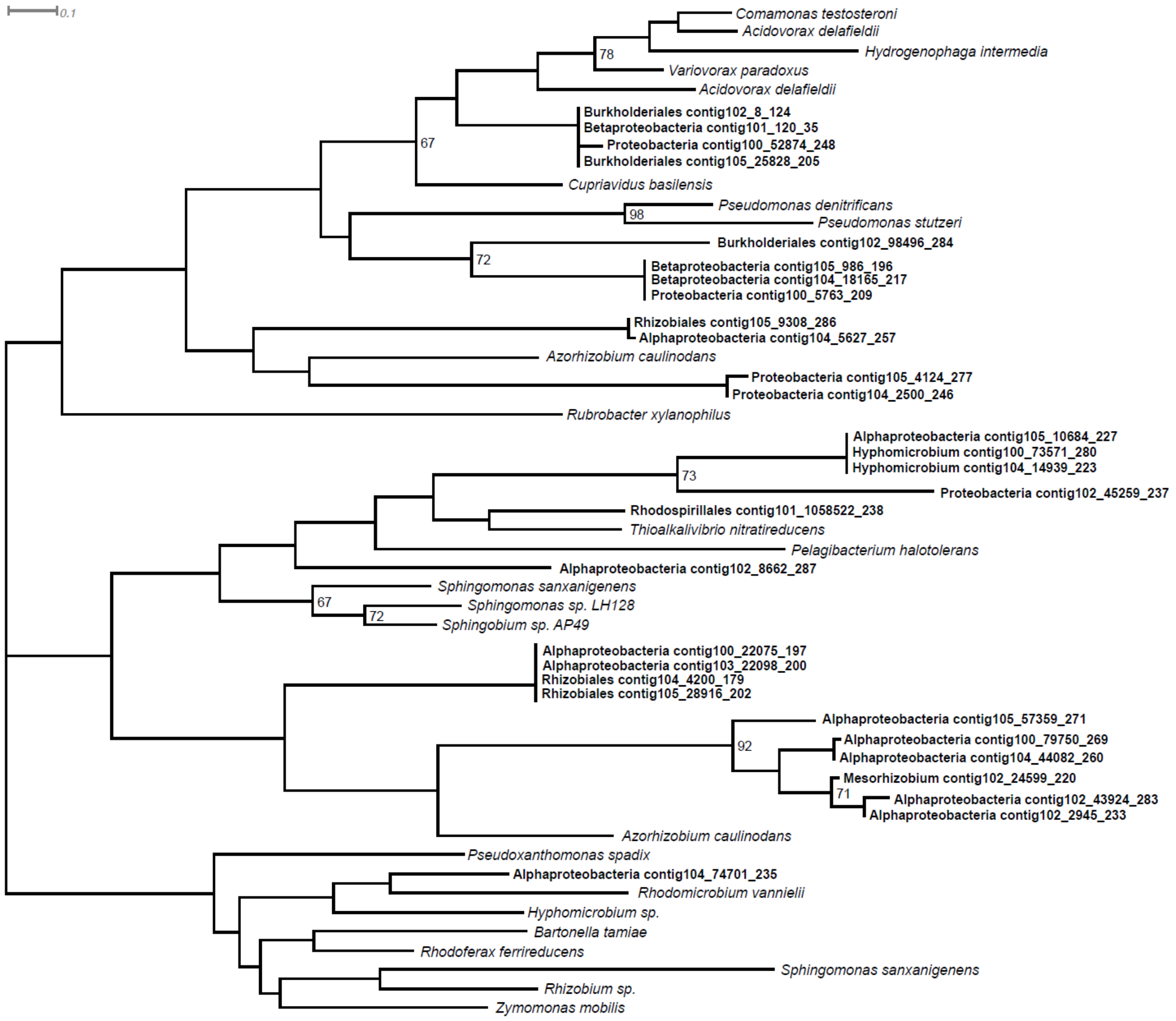


**Fig.S7** Phylogeny of putative heterotrophic *amoA* sequences retrieved from the metagenome and reference sequences obtained from (PF05145); the taxonomy of the metagenome-derived *amoA* sequences was inferred from the LCA of all of the genes on the respective contig.

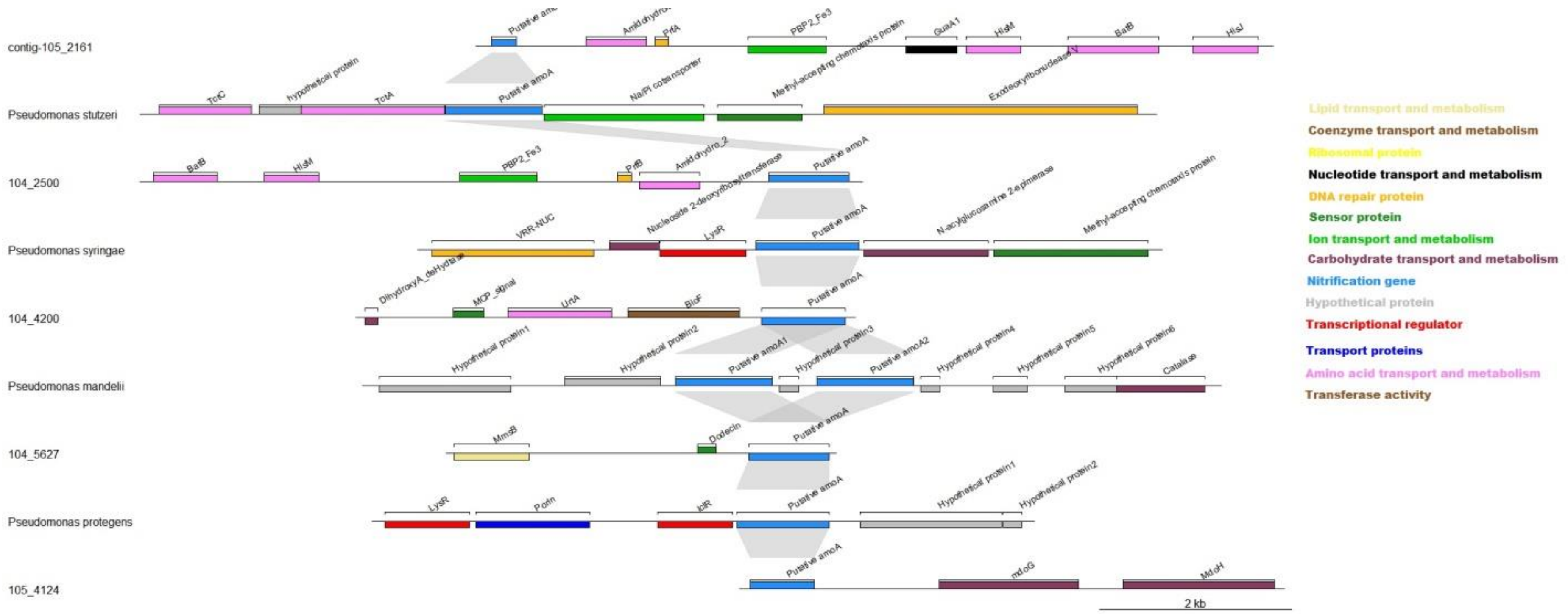


**Fig. S8** Gene synteny on some of the contigs on which the putative heterotrophic *amoA* sequences (PF 051459) were retrieved as well as gene synteny near reference sequences of *Pseudomonas* strains.

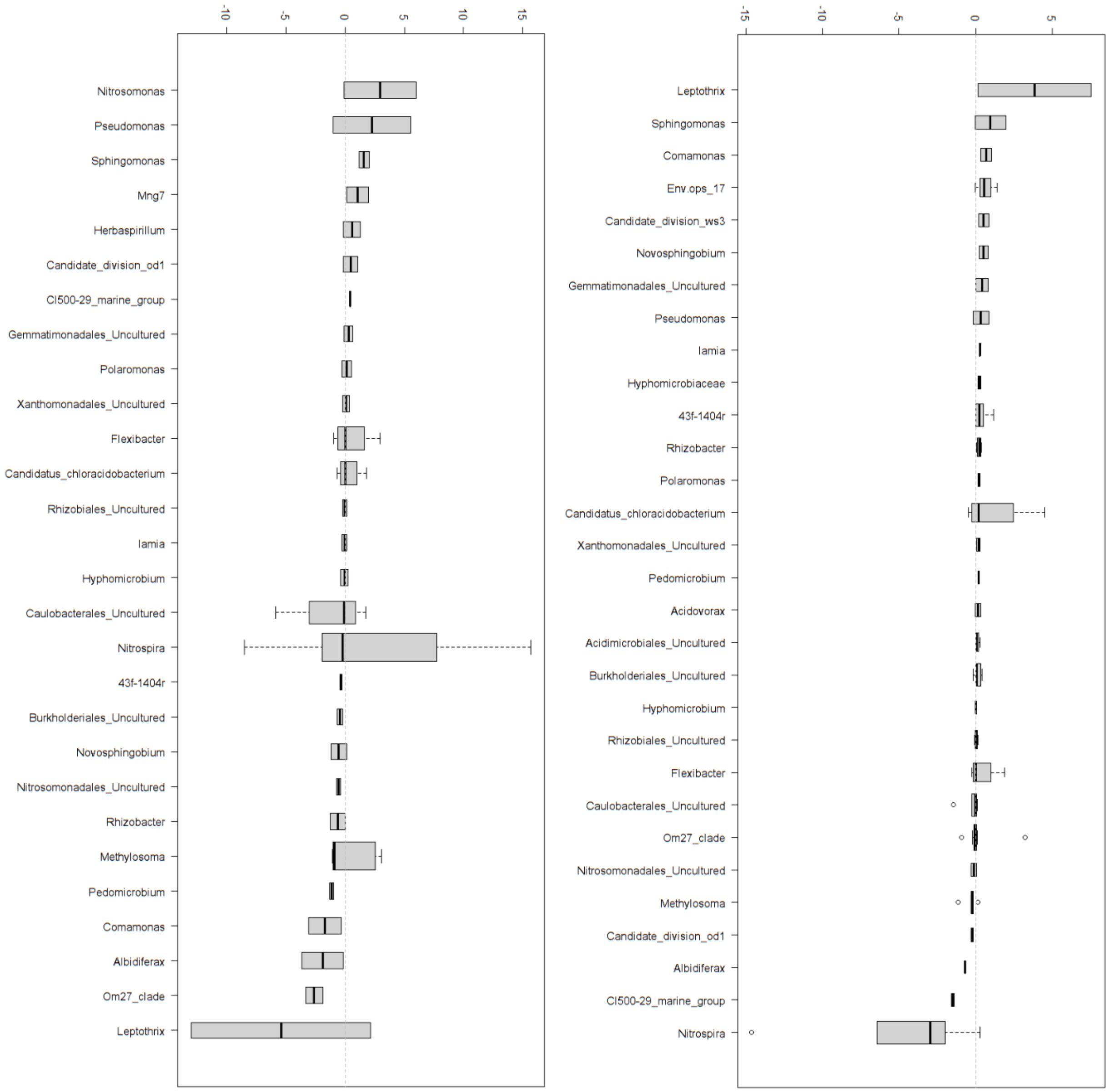


**Figure S9 A** Relative abundance shifts in 16S rDNA (left) and 16S rRNA (right) (in %) of the most dominant genera between day 0 and day 15 in columns 1 and 2.

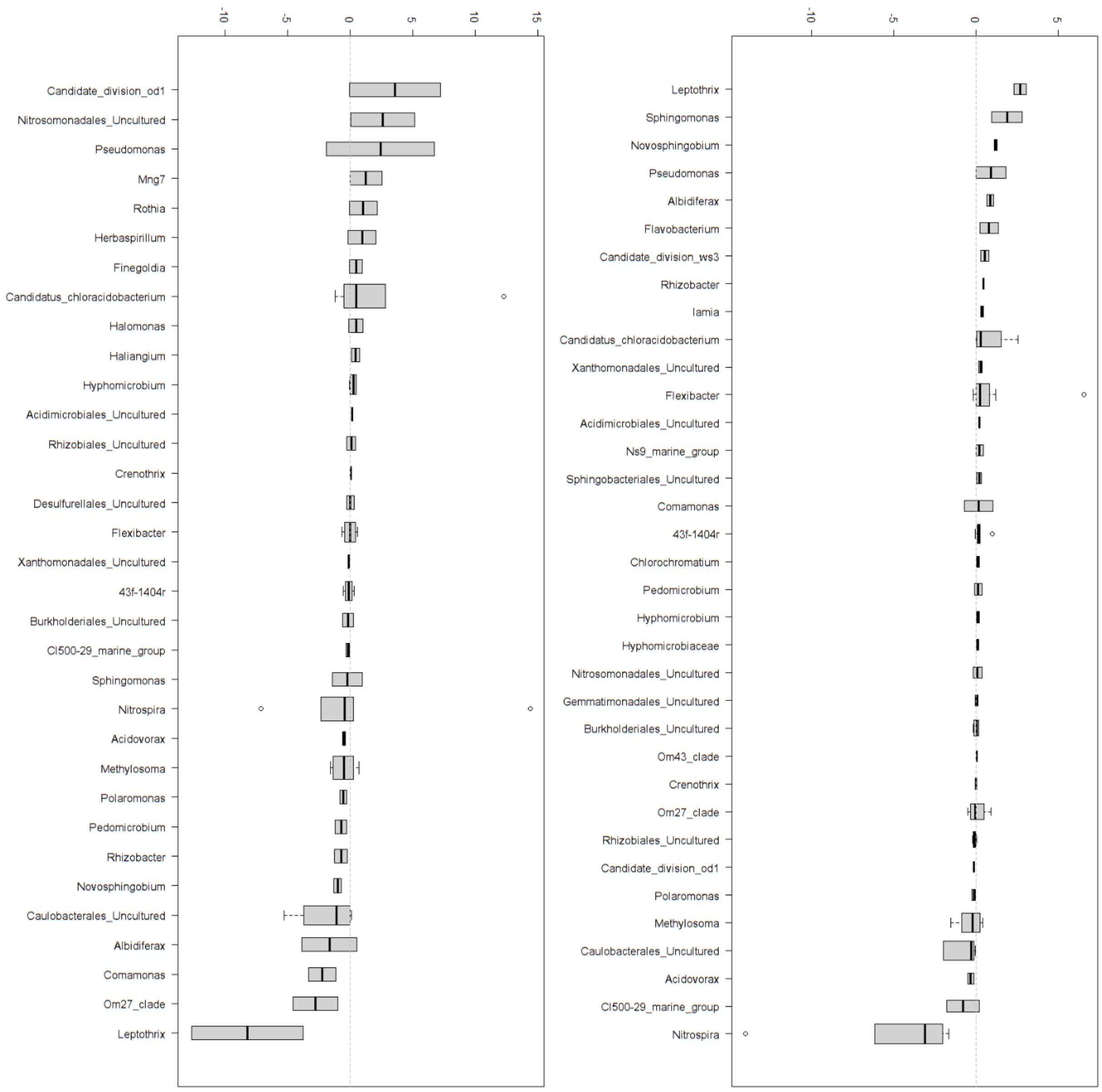


**Figure S9 B** Relative abundance shifts in 16S rDNA (left) and 16S rRNA (right) (in %) of the most dominant genera between day 0 and day 15 in columns 3 and 4.

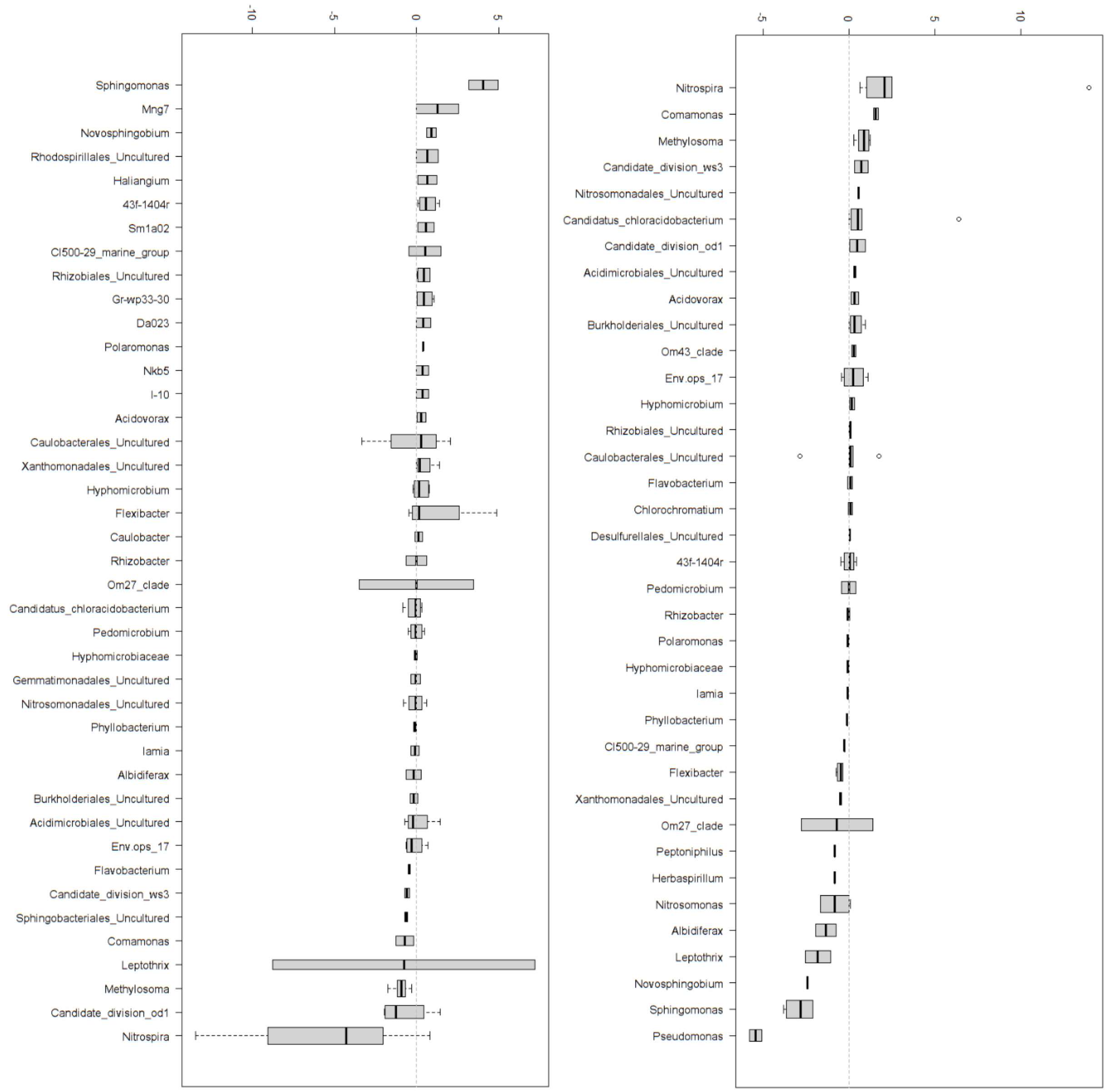


**Figure S9 C** Relative abundance shifts in 16S rDNA (left) and 16S rRNA (right) (in %) of the most dominant genera between day 0 and day 15 in columns 5 and 6.

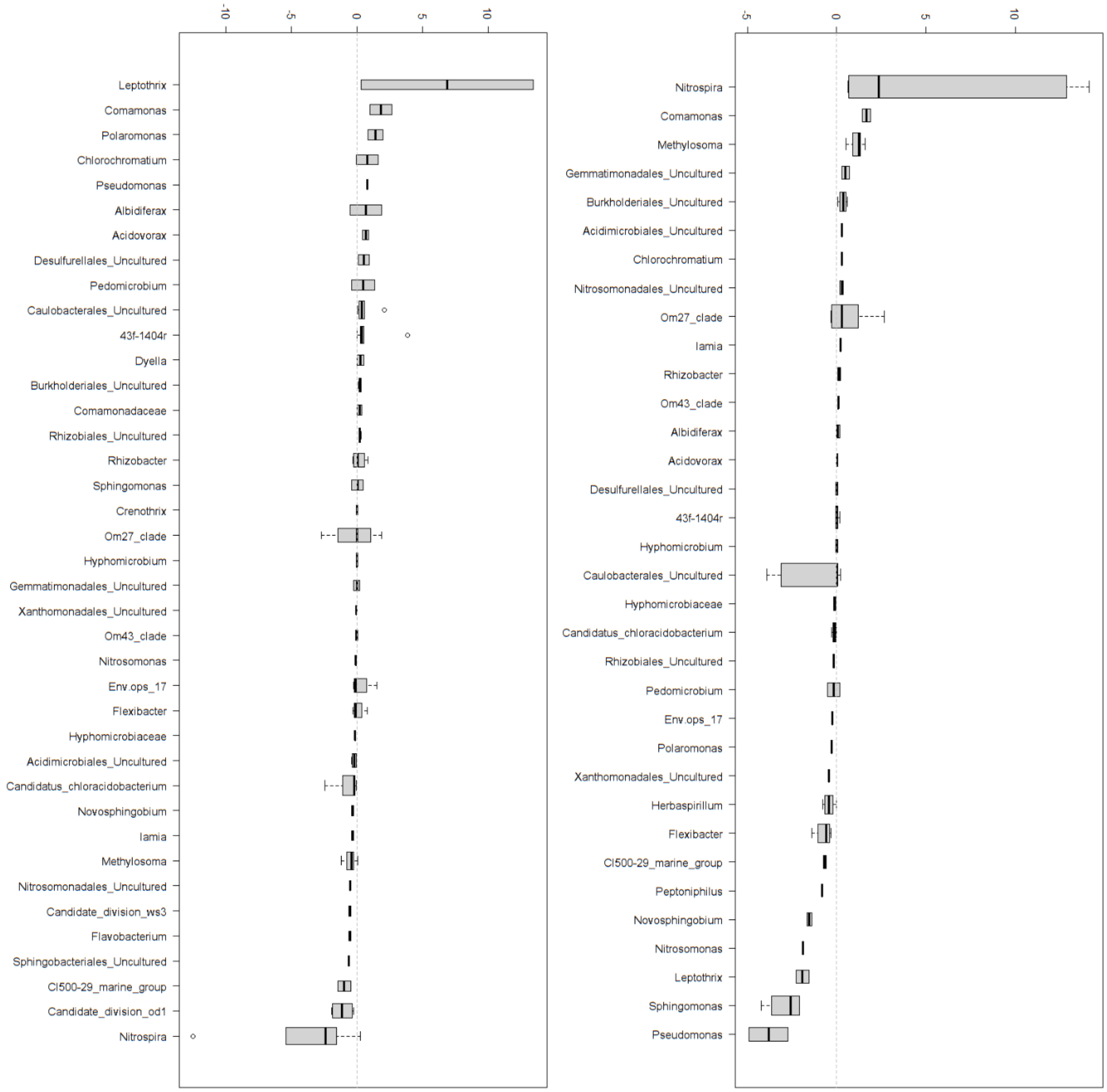


**Figure S9 D** Relative abundance shifts in 16S rDNA (left) and 16S rRNA (right) (in %) of the most dominant genera between day 0 and day 15 in columns 7 and 8.

### DNA and RNA-SIP reveal Nitrospira spp. as key drivers of nitrification in groundwater-fed biofilters

**Arda Gülay[1,4,*], Jane Fowler[1], Karolina Tatari, Bo Thamdrup[3], Hans-Jørgen Albrechtsen[1], Waleed Abu Al-Soud[2,] Søren J. Sørensen[2] and Barth F. Smets[1*]**

### Supplementary Materials & Methods

*Sampling sites and procedure*

Filter material samples were collected from an after-filter at Islevbro waterworks (Rødovre, Denmark). The waterworks was described in Gülay *et al.* (2) and influent and effluent water quality have been reported elsewhere (2, 3). Filter material samples were collected from three random horizontal locations of the after-filter using manual coring (12 cm inner diameter, 65 cm long). From the extracted filter material core, the top 10 cm was aseptically segregated on site and stored on ice for further use. A portion was frozen on-site in liquid nitrogen for RNA extractions.

*Column experiments and stable isotope labelling*

Column experiments were conducted using a continuous flow lab-scale system (3); columns were 2.6 cm in diameter 6 cm tall and filled with 5 cm (26.5 $cm^3$) of parent filter material. Effluent water from the investigated waterworks was used as the influent medium in all experiments to mimic full scale conditions. For nearly-complete isotope labeling of $HCO_3^-$,the total alkalinity in the effluent water was first removed by acidification followed by stripping, and then $^{13}C$-labeled, (or unlabeled, for control columns), bicarbonate ($HCO_3^-$) was added until the original alkalinity was reached.

The experimental design consisted of 4 treatments applied to labeled and unlabeled control columns. The experiments were organized in two phases of 4 columns each, and filter material was sampled just before the beginning of each experimental phase. The 4 treatments were applied by supplementing the influent water with: (i) 1 mg/L $NH_4^+$-N ($NH_4Cl$; Sigma-Aldrich, 254134), (ii) 1 mg/L $NH_4^+$-N and 100 µM ATU (N-Allylthiourea, Merck chemicals, 808158) , (iii) 1 mg/L $NO_2^-$-N ($NaNO_2$; Sigma-Aldrich, S2252), (iv) 1 mg/L $NO_2^-$-N and 1 mM ClO3$^-$ ($KClO_3^-$; 99%, Sigma-Aldrich, 12634 (Table 1). The flow rate was set to 40 ml/hr. The combination of flow rate and influent concentrations resulted in volumetric N loading rates (of $NH_4^+$-N or $NO_2^-$-N) that matched those experienced by the original full-scale filter (approx. 1.5 g N/$m^3$/hr) (3). We have previously observed that the $NH_4^+$-N loading rate (more than solely the $NH_4^+$-N influent concentration) is predictive of $NH_4^+$-N removal (3, 4).Test and control columns were operated for 15 days with continuous feeding. Preliminary calculations (assuming that all *Nitrospira* and *Nitrosomonas* cells in the columns were involved in ammonium oxidation, and a growth yield ranging from 0.03 to 0.01 mg biomass dry weight/mg ammonium N removed) indicated that the desired label incorporation (to 10% in all ammonium oxidizing cells) would require from 5 to 15 days incubation. In parallel columns fed with $NH_4^+$-N and $ClO_3^-$, the $ClO_3^-$ concentration was increased from 0.05 mM to 1 mM on the third day of operation. At the end of the runs, the columns were sacrificed and the filter material was immediately frozen in liquid nitrogen and stored at -80 °C for DNA and RNA extraction.

*Analytical methods*

Column effluents were sampled daily, filtered (20 µm), frozen and analyzed for $NH_4^+$ and $NO_2^-$ by colorimetric methods as described in Tatari et al. (3). Colorimetric analysis of ammonium in samples containing ATU was found to underestimate the $NH_4^+$ concentration (5) and thus $NH_4^+$ in these samples was quantified by flow injection analysis (6). $NO_3^-$ was quantified by Ion Chromatography (Dionex, ICS 1500 fitted with a guard column (Dionex, AG 22) and an analytical column (Dionex, ION PAC AS22). $NH_4^+$ removal (%) was calculated by subtracting effluent from influent $NH_4^+$ concentration and normalizing for the influent $NH_4^+$ concentration. Nitritation inhibition (%), was calculated as the difference in $NH_4^+$ removal of the control and the test columns. $NO_2^-$ removal (%) was calculated as the difference of effluent and produced $NO_2^-$ concentration, after correcting for trace $NO_2^-$ present in the water (0.016 mg/L $NO_2^-$) and normalization for the produced $NO_2^-$concentration. The $NO_2^-$produced by nitritation was calculated as the difference of influent and effluent $NH_4^+$ concentrations. Inhibition of nitratation (%), was calculated in the same way as the difference of $NO_2^-$ removal between the control (Col.2, Col.3, Col.6 and Col.7) and test columns (Col.1, Col.4, Col.5 and Col.8). $NO_3^-$ accumulation (%) was calculated by subtracting effluent from influent $NO_3^-$ concentration.

Eventual losses of N were checked in the system by comparing the total influent $NH_4^+$, $NO_2^-$ and $NO_3^-$ with the total N concentration of the same species each day of the experiment. Comparison was applied by a 2-tailed t-test, setting a significance level of 0.05 and once normal distribution of the data was checked. This analysis was done in treatments where ATU was not added, due to interference of the inhibitor with the analytical method used for $NH_4^+$quantification(7). In these columns, $NH_4^+$was re-measured with flow-injection, but due to the different analytical methods used, no N balance check was done. In all other columns, the check showed that total influent and effluent N fitted, except from one case where N loss was observed. N balances were calculated by using the Equation 1:

$$\Delta \mathrm{N} = [\mathrm{N}-\mathrm{NH_4^+} + \mathrm{N}-\mathrm{NO_2^-} + \mathrm{N}-\mathrm{NO_3^-}]_{\mathrm{in}} - [\mathrm{N}-\mathrm{NH_4^+} + \mathrm{N}-\mathrm{NO_2^-} + \mathrm{N}-\mathrm{NO_3^-}]_{\mathrm{eff}} \qquad \text{Eq.1}$$

$$*\,\sigma = \sqrt{2\sigma_{NH4}^2 + 2\sigma_{NO2}^2 + 2\sigma_{NO3}^2}$$

*Standard deviation of each species based on steady-state influent values

*RNA-DNA extraction and stable isotope probing (SIP)*

Filter material samples collected from the full scale filter and the sacrificed columns were subject to DNA and RNA extraction. Genomic DNA was extracted from 0.5 g of drained filter material using the MP FastDNA™ SPIN Kit (MP Biomedicals LLC., Solon, USA) according to manufacturer's instructions. The concentration and purity of extracted DNA were checked by spectrophotometry (NanoDrop Technologies, Wilmington, DE, USA). RNA was extracted from frozen filter material samples (-80 °C) with a MoBio PowerSoil Total RNA Isolation Kit (#12866-25) according to manufacturer's instructions. The RNA was further purified with a Qiagen AllPrep DNA/RNA Mini Kit (Hilden, Germany) and quantified with a Ribogreen RNA-quantification kit (Invitrogen, Eugene, OR, USA). 650 ng of purified RNA were fractionated after density gradient ultracentrifugation at 38400 rpm for 72 h at 20 °C (8). RNA purification in each fraction was performed according to Whiteley et al. (9). The concentration of purified RNA was determined using a Ribogreen RNA-quantification kit.

Density gradient ultracentrifugation of DNA isolated from columns and full-scale was performed according to Neufeld et al. (2007). Briefly, 1.6 μg of DNA in CsCl with a final density of approximately 1.725 g/mL was subject to ultracentrifugation at 44800 rpm for 44 h, 20°C in a ultracentrifuge (Beckmann) with a VTi65.2 rotor (Beckmann). Gradients were fractionated into 250 μL fractions, density was determined by refractometry and DNA was recovered by precipitation with PEG. DNA concentration was determined using a Picogreen high sensitivity dsDNA quantification kit (Invitrogen).

*PCR amplification and tag sequencing*

RNA purified from density gradient fractions, sacrificed column samples and full scale filter samples were reverse transcribed using reverse primer 1492R. 10 ng of cDNA and DNA from these samples (Table 1) were used to amplify the V3-V4 regions of bacterial 16S rRNA genes using the Phusion (Pfu) DNA polymerase (Finnzymes, Finland) and 16S rRNA gene targeted (rDNA) modified universal primers PRK341F and PRK806R (10). PCR was performed as described in Gülay et al. (11). Pyrosequencing was applied in a two-region 454 run on a 70-75 GS PicoTiterPlate using a Titanium kit and GS FLX pyrosequencing system at the National High-throughput DNA Sequencing Center (Copenhagen, DK). Purified DNA from all fractions was amplified as described above and sequenced on an Illumina MiSeq platform at the National High-throughput DNA Sequencing Center (Copenhagen, DK).

*Bioinformatic analysis*

Raw sequence 454 data from RNA-SIP samples were quality-checked (denoised) with Ampliconnoise (12) and chimeras were removed with UCHIME (13) using default settings. Raw sequence Miseq Illumina data from DNA-SIP samples were quality-checked with mothur and chimeras were removed with UCHIME (13) using a reference dataset. Sequence libraries were combined and trimmed to 418 bp. All analyses were performed in QIIME 1.9.1 (14). High quality sequences were clustered into OTUs at 99% pairwise identity using UCLUST (15) in de novo mode with default settings. Representative sequences from each OTU were aligned against the curated silva.seed_v123.align database. Taxonomic assignment of OTUs was implemented using the BLAST algorithm (16) against the Silva128 database (17). Sequences with less than 90% similarity to reference sequences were deemed unclassified.

**Filter 1** The absolute mass of each OTU was estimated by multiplying the mass of DNA or RNA in each SIP fraction with relative abundance associated with each OTU in the respective amplicon library. R codes related to this conversion and other operations can be found in https://github.com/ardagulay. Labelled OTUs in DNA-SIP were detected by comparing the buoyant density of each OTU in replicate columns fed with $H^{13}CO_3^-$ and $H^{12}CO_3^-$ using Eq.2. The ratio of the weighted buoyant density of each OTU was calculated based on its concentration in all fractions and buoyant density of these fractions:

$$\mu_{OTU\_i} = \frac{Y_{LF1}*X_{LF1}+Y_{LF2}*X_{LF2}+\cdots\ldots+Y_{LFn}*X_{LFn}}{Y_{UF1}*X_{UF1}+Y_{UF2}*X_{UF2}+\cdots\ldots+Y_{UFn}*X_{UFn}} \qquad \text{Eq.2}$$

where $\mu_{OTU\_i}$ is the ratio of the weighted buoyant density of the i-th OTU across the gradient, $Y_{LF}$ and $Y_{UF}$ are the OTU's mass (ng DNA) in each fraction of the $H^{13}CO_3^-$ fed and $H^{12}CO_3^-$ columns respectively, $X_{LF}$ and $X_{UF}$ are the densities of each fraction in the $H^{13}CO_3^-$ fed and $H^{12}CO_3^-$ column respectively. OTUs for which the ratio of the weighted buoyant density was higher than 1 were selected as $^{13}C$ labelled via DNA-SIP during the specific treatment.

A similar reasoning was applied to detect OTUs labelled via RNA-SIP in a specific treatment As only selected fractions were subject to amplicon sequencing, the mean buoyant density of each OTU in the $H^{13}CO_3^-$ fed) and $H^{12}CO_3^-$ column were calculated as described in Zemb et al (2012). Assuming that RNA concentrations of OTUs follow a normal distribution (Fig. S4) across the gradient, the mean buoyant density of the i-th OTU can be calculated according to Eq.(3):

$$\mu_{OTU\_i} = \frac{2\sigma^2_{total\_UN\_RNA} * ln\left(\frac{Y_{LL}}{Y_{HL}}\right) - x^2_{HL} + x^2_{LL}}{2(x_{HL}+x_{LL})} \qquad \text{Eq.3}$$

where $\sigma_{total\_UN\_RNA}$ is the standard deviation of the RNA distribution derived from the RNA (ng) in the gradient (Fig S3-S4) in the $H^{12}CO_3^-$ fed colum, $x_{HL}$ and $x_{LL}$ are the densities of the representative heavy and light fractions of the $H^{13}CO_3^-$ fed column respectively, and $y_{LL}$ and $y_{HL}$ are the mass of a specific OTUs (ng RNA) in the LL (LL;<1.80 CsTFA buoyant density) and HL (HL>1.80 CsTFA buoyant density) fractions respectively. Among the sequenced fractions, light and heavy fractions with the highest RNA mass were selected. The mean buoyant density of the i-th OTU was calculated for both $H^{12}CO_3^-$ fed and $H^{13}CO_3^-$ fed column. Buoyant density shifts of the i-th OTU were calculated as the difference in the calculated mean buoyant densities between the replicate columns. OTU with buoyant density shifts higher than zero were selected as $^{13}C$ labelled in RNA-SIP in a specific treatment. R codes related to the detection of labelled OTUs in DNA and RNA –SIP can be found in https://github.com/ardagulay.

Taxonomic assignment of the selected OTUs was implemented using the BLAST algorithm (16) against the Silva128 database (17) after re-alignment with web-based SINA v1.2.11 (18).

**Filter 2** Genera which contained a minimum of 10 labelled OTUs in both RNA and DNA-SIP were selected as labelled genera; OTUs not belonging to the selected genera were deemed unlabelled and excluded from further analysis.

**Filter 3** We then used bootstrap resampling (with replacement, 1,000 iterations) of replicates (labelled OTUs) within each detected genus to estimate genus-specific 90% CIs for buoyant density changes in both DNA and RNA (Table S1). For each bootstrap iteration replicates (with replacement) equal to the number of labelled OTUs within the selected genus were drawn and OTUs below genus-specific 90% CIs were excluded from the analysis. R codes related to the bootstrap iteration can be found in https://github.com/ardagulay.

**Filter 4** To identify ammonium and nitrite oxidizing phylotypes, we further examined the labelled OTUs within each treatment and excluded all OTUs with buoyant density shift values lower than the maximum buoyant density shift value of labelled *Nitrosomonas* and *Nitrospira* OTUs, respectively.

**Filter 5** We, then, selected the genera which contained OTUs in both RNA and DNA-SIP; OTUs not belonging to the selected genera were excluded from further analysis.

**Filter 6** We, subsequently, compared the OTUs between treatments according to the scheme in Fig.4.a. to identify the ammonia and nitrite oxidizing phylotypes: only OTUs that were exclusively present in the solely $NH_4^+$ treatment (i.e. were absent from $NH_4^+$ + ATU, $NH_4^+$ + $ClO_3^-$, and $NO_2^-$

treatments), were retained as ammonia oxidizing phylotypes. Similarly, only OTUs that were exclusively present in the $NO_2^-$ treatment and were absent from the $NH_4^+$ + Chlorate treatment were retained as nitrite oxidizing phylotypes.

Finally, the retained ammonia and nitrite oxidizing phylotypes were compared to the genera that displayed relative DNA and RNA abundance shifts (calculated from total DNA and RNA sequence libraries retrieved from the samples taken at the start and end of the experiment) higher than what was observed for phylotypes of the *Nitrosomonas* and *Nitrospira* genus, respectively.

Phylogenetic analysis of the labelled OTUs was implemented with Fast Tree (19) in QIIME and iTOL was used for visualization (Letunic and Bork, 2007; http://itol.embl.de/). Label percentage of OTUs was calculated from buoyant density shifts; the estimated buoyant DNA and RNA shift for each OTU was divided by the total observed shift of labelled OTUs for DNA (Eqn 4) and RNA SIP (Eqn 5), respectively.

$$\mu_{OTU\_i\_\%\_DNA} = \frac{\mu_{OTU\_i}}{\mu_{OTU\_1} + \mu_{OTU\_2} \cdots\cdots + \mu_{OTU\_n}} * 100 \qquad \text{Eq 4}$$

$$\mu_{OTU\_i\_\%\_RNA} = \frac{\mu_{OTU\_i}}{\mu_{OTU\_1} + \mu_{OTU\_2} \cdots\cdots + \mu_{OTU\_n}} * 100 \qquad \text{Eq 5}$$

Changes in DNA and RNA relative abundance were calculated (for all OTUs identified as labeled) from the non-fractionated DNA and RNA sequence libraries retrieved from the samples taken at the onset and after 15 days of column operation. Abundance data in samples from replicate (labelled and unlabelled) columns were averaged to obtain final abundances. Pairwise comparisons of OTUs between 0 and 15 days of operation was implemented using total DNA and RNA based 16S rRNA libraries after subsampling the total sequence pools to 2500.

The metagenome of the parent full-scale filter microbial community has been previously described (21). The ammonia monooxygenase subunit A (*amoA*) genes annotated as PF05145 (heterotrophic *amoA*) were extracted from the metagenome. Amino acid sequences were aligned with reference sequences using MUSCLE (22) and maximum likelihood trees were constructed in MEGA7.

nonylphenol degraders in activated sludge. Water Res 46:601–610.

2. Gülay A, Musovic S, Albrechtsen H-JH-JH-J, Smets BF. 2013. Neutrophilic iron-oxidizing bacteria: occurrence and relevance in biological drinking water treatment. Water Sci Technol Water Supply 13:1295.
3. Tatari K, Smets BF, Albrechtsen HJ. 2013. A novel bench-scale column assay to investigate site-specific nitrification biokinetics in biological rapid sand filters. Water Res 47:6380–6387.
4. Lee CO, Boe-Hansen R, Musovic S, Smets B, Albrechtsen H-J, Binning P. 2014. Effects of dynamic operating conditions on nitrification in biological rapid sand filters for drinking water treatment. Water Res 64:226–236.
5. Gülay A, Tatari K, Musovic S, Mateiu R V., Albrechtsen H-J, Smets BF. 2014. Internal porosity of mineral coating supports microbial activity in rapid sand filters for groundwater treatment. Appl Environ Microbiol 80:7010–20.
6. Hall P, Aller R. 1992. Rapid, small-volume, flow-injection analysis for sigma-CO2 and NH4+ in marine and fresh-waters. Limnol Oceanogr 37:1113–1119.
7. Tatari K, Gülay A, Thamdrup B, Albrechtsen HJ, Smets BF. 2017. Challenges in using allylthiourea and chlorate as specific nitrification inhibitors. Chemosphere 182:301–305.
8. Vandieken V, Pester M, Finke N, Hyun J-H, Friedrich MW, Loy A, Thamdrup B. 2012. Three manganese oxide-rich marine sediments harbor similar communities of acetate-oxidizing manganese-reducing bacteria. ISME J 6:2078–90.
9. Whiteley AS, Thomson B, Lueders T, Manefield M. 2007. RNA stable-isotope probing. Nat Protoc 2:838–844.
10. Yu Y, Lee C, Kim J, Hwang S. 2005. Group-specific primer and probe sets to detect methanogenic communities using quantitative real-time polymerase chain reaction. Biotechnol Bioeng 89:670–9.
11. Gülay A, Musovic S, Albrechtsen H-J, Al-Soud WA, Sørensen SJ, Smets BF. 2016. Ecological patterns, diversity and core taxa of microbial communities in groundwater-fed rapid gravity filters. ISME J 10:2209–2222.
12. Quince C, Lanzen A, Davenport RJ, Turnbaugh PJ. 2011. Removing noise from pyrosequenced amplicons. BMC Bioinformatics 12:38.
13. Edgar RC, Haas BJ, Clemente JC, Quince C, Knight R. 2011. UCHIME improves sensitivity and speed of chimera detection. Bioinformatics 27:2194–200.
14. Caporaso JG, Kuczynski J, Stombaugh J, Bittinger K, Bushman FD, Costello EK, Fierer N, Peña AG, Goodrich JK, Gordon JI, Huttley GA, Kelley ST, Knights D, Koenig JE, Ley RE,

Lozupone CA, Mcdonald D, Muegge BD, Pirrung M, Reeder J, Sevinsky JR, Turnbaugh PJ, Walters WA, Widmann J, Yatsunenko T, Zaneveld J, Knight R. 2010. QIIME allows analysis of high- throughput community sequencing data. Nat Methods 7:335–336.

15. Edgar RC. 2010. Search and clustering orders of magnitude faster than BLAST. Bioinformatics 26:2460–1.

16. Altschul SF, Gish W, Miller W, Myers EW, J. LD. 1990. Basic local alignment search tool. J Mol Biol 215.

### DNA and RNA-SIP reveal Nitrospira spp. as key drivers of nitrification in groundwater-fed biofilters

**Arda Gülay[1,4,*], Jane Fowler[1], Karolina Tatari, Bo Thamdrup[3], Hans-Jørgen Albrechtsen[1], Waleed Abu Al-Soud[2,] Søren J. Sørensen[2] and Barth F. Smets[1*]**

### Supplementary Tables

**Table S1** Genus specific 90% CIs for the change in DNA buoyant density

| Comparison | Taxa | 90%CI |
|---|---|---|
| **DNA_C1vsC2** | **Nitrospira** | **27.29490368** |
| DNA_C1vsC2 | Hyphomicrobium | 6.965868301 |
| DNA_C1vsC2 | OM27_clade | 12.252273 |
| DNA_C1vsC2 | Blastocatella | 4.527497928 |
| DNA_C1vsC2 | Sphingomonas | 7.832605989 |
| DNA_C1vsC2 | Methyloglobulus | 9.972687698 |
| DNA_C1vsC2 | Woodsholea | 11.51530212 |
| DNA_C1vsC2 | uncultured_Latescibacteria_bacterium | 4.395130052 |
| DNA_C1vsC2 | Pseudomonas | 20.78427096 |
| DNA_C1vsC2 | Variovorax | 3.703893218 |
| DNA_C1vsC2 | Nitrosococcus | 2.560913888 |
| DNA_C1vsC2 | Pedomicrobium | 7.954590866 |
| DNA_C1vsC2 | uncultured | 4.788737221 |
| DNA_C1vsC2 | ABS-19 | 6.385744045 |
| DNA_C1vsC2 | Nitrosomonas | 2.349722921 |
| DNA_C1vsC2 | Rhizobacter | 5.251468261 |
| DNA_C1vsC2 | CL500-29_marine_group | 5.775774936 |
| DNA_C1vsC2 | Acidovorax | 3.243152628 |
| DNA_C4vsC3 | Woodsholea | 8.746445583 |
| DNA_C4vsC3 | uncultured_Latescibacteria_bacterium | 6.922797297 |
| DNA_C4vsC3 | Blastocatella | 5.900721972 |
| DNA_C4vsC3 | ABS_19 | 4.864921525 |
| DNA_C4vsC3 | Sphingomonas | 8.427538926 |
| DNA_C4vsC3 | Azospira | 3.467360657 |
| DNA_C4vsC3 | Pedomicrobium | 11.24330224 |
| DNA_C4vsC3 | Hyphomicrobium | 11.25563077 |
| DNA_C4vsC3 | Pseudomonas | 21.77335039 |
| DNA_C4vsC3 | Nitrospira | 8.999803853 |
| DNA_C4vsC3 | Nitrosomonas | 5.042264724 |
| DNA_C4vsC3 | Nitrosococcus | 2.410856048 |
| DNA_C4vsC3 | Methyloglobulus | 7.6875397 |
| DNA_C4vsC3 | CL500-29_marine_group | 10.03613588 |
| DNA_C4vsC3 | Acidovorax | 6.612988123 |
| DNA_C5vsC6 | Woodsholea | 34.61995489 |
| DNA_C5vsC6 | uncultured_Latescibacteria_bacterium | 12.06266939 |
| DNA_C5vsC6 | uncultured | 7.157708609 |
| DNA_C5vsC6 | Sphingomonas | 11.87161387 |
| DNA_C5vsC6 | Pedomicrobium | 27.32630925 |
| DNA_C5vsC6 | OM27_clade | 35.22452854 |
| DNA_C5vsC6 | Nitrospira | 69.06265794 |
| DNA_C5vsC6 | Nitrosococcus | 6.604486979 |
| DNA_C5vsC6 | Methyloglobulus | 21.34205684 |
| DNA_C5vsC6 | Hyphomicrobium | 27.3900279 |
| DNA_C5vsC6 | CL500-29_marine_group | 21.98392165 |

| | | |
|---|---|---|
| DNA_C5vsC6 | Blastocatella | 14.69858365 |
| DNA_C8vsC7 | Woodsholea | 19.99953085 |
| DNA_C8vsC7 | OM27_clade | 21.00440171 |

**Table S2** Genus specific 90% CIs for the change in RNA buoyant density

| Comparison | Taxa | 90%CI |
|---|---|---|
| RNA.C1vsC2 | Variovorax | 0.010946202 |
| RNA.C1vsC2 | Methyloglobulus | 0.012084529 |
| RNA.C1vsC2 | ABS-19 | 0.009321392 |
| RNA.C1vsC2 | CL500-29_marine_group | 0.013380092 |
| RNA.C1vsC2 | Acidovorax | 0.010681521 |
| RNA.C1vsC2 | OM27_clade | 0.012539093 |
| RNA.C1vsC2 | Nitrosococcus | 0.010281794 |
| RNA.C1vsC2 | Sphingomonas | 0.011672409 |
| RNA.C1vsC2 | Rhizobacter | 0.010244547 |
| RNA.C1vsC2 | Woodsholea | 0.011844113 |
| RNA.C1vsC2 | Pseudomonas | 0.010696879 |
| RNA.C1vsC2 | uncultured | 0.010812986 |
| RNA.C1vsC2 | Nitrospira | 0.013709626 |
| RNA.C1vsC2 | Blastocatella | 0.012385466 |
| RNA.C1vsC2 | uncultured_Latescibacteria_bacterium | 0.009498014 |
| RNA.C1vsC2 | Nitrosomonas | 0.009555628 |
| RNA.C1vsC2 | Pedomicrobium | 0.011050473 |
| RNA.C1vsC2 | Hyphomicrobium | 0.010592564 |
| RNA.C4vsC3 | Woodsholea | 0.008621579 |
| RNA.C4vsC3 | uncultured_Latescibacteria_bacterium | 0.007414314 |
| RNA.C4vsC3 | Blastocatella | 0.00661408 |
| RNA.C4vsC3 | ABS_19 | 0.008422849 |
| RNA.C4vsC3 | Sphingomonas | 0.007734611 |
| RNA.C4vsC3 | Azospira | 0.006711897 |
| RNA.C4vsC3 | Pedomicrobium | 0.007508337 |
| RNA.C4vsC3 | Hyphomicrobium | 0.00721053 |
| RNA.C4vsC3 | Pseudomonas | 0.009573882 |
| RNA.C4vsC3 | Nitrospira | 0.008457103 |
| RNA.C4vsC3 | Nitrosomonas | 0.006827565 |
| RNA.C4vsC3 | Nitrosococcus | 0.006299523 |
| RNA.C4vsC3 | Methyloglobulus | 0.006913547 |
| RNA.C4vsC3 | CL500-29_marine_group | 0.006162106 |
| RNA.C4vsC3 | Acidovorax | 0.008380405 |
| RNA.C5vsC6 | Woodsholea | 0.023065677 |
| RNA.C5vsC6 | uncultured_Latescibacteria_bacterium | 0.022863257 |
| RNA.C5vsC6 | uncultured | 0.021014205 |
| RNA.C5vsC6 | Sphingomonas | 0.022086214 |
| RNA.C5vsC6 | Pedomicrobium | 0.021504796 |
| RNA.C5vsC6 | OM27_clade | 0.027101066 |
| RNA.C5vsC6 | Nitrospira | 0.023046592 |
| RNA.C5vsC6 | Nitrosococcus | 0.02328091 |
| RNA.C5vsC6 | Methyloglobulus | 0.02270309 |
| RNA.C5vsC6 | Hyphomicrobium | 0.020008472 |
| RNA.C5vsC6 | CL500-29_marine_group | 0.022994518 |

| | | |
|---|---|---|
| RNA.C5vsC6 | Blastocatella | 0.021751132 |
| RNA.C8vsC7 | Woodsholea | 0.000814697 |
| RNA.C8vsC7 | OM27_clade | 0.001081158 |

**Table S3** Blast hits to the putative amoA sequences

| Query | Description | Accession | e-value | score |
|---|---|---|---|---|
| **gene_274\|GeneMark.hmm\|128_aa\|-\|100\|486** | hypothetical protein [Methyloferula stellata] | gi\|519017702\|ref\|WP_020173577.1\| | 2.49E-49 | 169 |
| **gene_238\|GeneMark.hmm\|188_aa\|+\|2\|565** | hypothetical protein [Tistlia consotensis] | gi\|1184553655\|ref\|WP_085121045.1\| | 2.01E-59 | 197 |
| **gene_284\|GeneMark.hmm\|159_aa\|-\|152\|631** | membrane protein [Cupriavidus sp. amp6] | gi\|656005008\|ref\|WP_029046450.1\| | 1.20E-37 | 140 |
| **gene_269\|GeneMark.hmm\|240_aa\|-\|1\|720** | putative ammonia monooxygenase [Azospirillum brasilense] | gi\|504008002\|ref\|WP_014241996.1\| | 4.71E-43 | 157 |
| **gene_280\|GeneMark.hmm\|253_aa\|+\|1\|759** | hypothetical protein [Hyphomicrobium sp. CS1BSMeth3] | gi\|1119410285\|ref\|WP_072385832.1\| | 6.52E-72 | 232 |
| **gene_235\|GeneMark.hmm\|197_aa\|+\|1\|594** | ammonia monooxygenase [Phyllobacterium sp. YR531] | gi\|495398156\|ref\|WP_008122856.1\| | 2.22E-66 | 215 |
| **gene_248\|GeneMark.hmm\|175_aa\|-\|418\|945** | hypothetical protein [Ramlibacter sp. Leaf400] | gi\|946973621\|ref\|WP_055894610.1\| | 5.07E-52 | 177 |
| **gene_271\|GeneMark.hmm\|262_aa\|-\|1\|786** | hypothetical protein [Azospirillum brasilense] | gi\|916533225\|ref\|WP_051140667.1\| | 4.79E-47 | 168 |
| **gene_237\|GeneMark.hmm\|334_aa\|-\|3\|1004** | hypothetical protein [Hyphomicrobium sp. CS1BSMeth3] | gi\|1119410285\|ref\|WP_072385832.1\| | 4.89E-89 | 279 |
| **gene_260\|GeneMark.hmm\|238_aa\|+\|2\|718** | ammonia monooxygenase [Labrenzia alba] | gi\|944196777\|ref\|WP_055678338.1\| | 6.67E-46 | 164 |
| **gene_283\|GeneMark.hmm\|227_aa\|+\|399\|1079** | hypothetical protein [Azospirillum brasilense] | gi\|916533225\|ref\|WP_051140667.1\| | 2.28E-43 | 157 |
| **gene_202\|GeneMark.hmm\|328_aa\|+\|575\|1558** | hypothetical protein [Rhodoplanes sp. Z2-YC6860] | gi\|1056597137\|ref\|WP_068028260.1\| | 4.65E-103 | 315 |
| **gene_220\|GeneMark.hmm\|350_aa\|-\|179\|1231** | ammonia monooxygenase [Polymorphum gilvum] | gi\|503419284\|ref\|WP_013653945.1\| | 7.83E-69 | 227 |
| **gene_205\|GeneMark.hmm\|291_aa\|+\|809\|1681** | hypothetical protein [Caldimonas taiwanensis] | gi\|1180938568\|ref\|WP_084362242.1\| | 6.46E-80 | 254 |
| **gene_200\|GeneMark.hmm\|368_aa\|-\|567\|1673** | hypothetical protein [Rhodoplanes sp. Z2-YC6860] | gi\|1056597137\|ref\|WP_068028260.1\| | 6.38E-105 | 321 |
| **gene_197\|GeneMark.hmm\|368_aa\|+\|44\|1150** | hypothetical protein [Rhodoplanes sp. Z2-YC6860] | gi\|1056597137\|ref\|WP_068028260.1\| | 6.38E-105 | 321 |
| **gene_217\|GeneMark.hmm\|311_aa\|-\|636\|1571** | membrane protein [Pseudogulbenkiania sp. MAI-1] | gi\|635657271\|ref\|WP_024304138.1\| | 1.23E-75 | 243 |
| **gene_223\|GeneMark.hmm\|380_aa\|-\|341\|1483** | hypothetical protein [Hyphomicrobium sp. CS1BSMeth3] | gi\|1119410285\|ref\|WP_072385832.1\| | 3.60E-127 | 378 |
| **gene_227\|GeneMark.hmm\|366_aa\|-\|1878\|2978** | hypothetical protein [Hyphomicrobium sp. CS1BSMeth3] | gi\|1119410285\|ref\|WP_072385832.1\| | 7.66E-127 | 377 |
| **gene_287\|GeneMark.hmm\|248_aa\|-\|2178\|2924** | hypothetical protein [Rhizobiales bacterium CCH3-A5] | gi\|1177645899\|ref\|WP_082736595.1\| | 2.94E-80 | 253 |
| **gene_268\|GeneMark.hmm\|287_aa\|-\|2379\|3242** | ammonia monooxygenase [Rhizobium sp. NT-26] | gi\|918947167\|ref\|WP_052641815.1\| | 3.31E-36 | 140 |
| **gene_209\|GeneMark.hmm\|352_aa\|+\|2469\|3527** | membrane protein [Pseudogulbenkiania sp. MAI-1] | gi\|635657271\|ref\|WP_024304138.1\| | 1.14E-83 | 265 |
| **gene_257\|GeneMark.hmm\|292_aa\|-\|3418\|4296** | ammonia monooxygenase [Rhizobium sp. NT-26] | gi\|918947167\|ref\|WP_052641815.1\| | 9.85E-38 | 144 |
| **gene_179\|GeneMark.hmm\|368_aa\|-\|4069\|5175** | hypothetical protein [Rhodoplanes sp. Z2-YC6860] | gi\|1056597137\|ref\|WP_068028260.1\| | 8.96E-108 | 328 |
| **gene_277\|GeneMark.hmm\|271_aa\|+\|1\|816** | AbrB family transcriptional regulator [Thauera sp. 27] | gi\|489025714\|ref\|WP_002936124.1\| | 3.73E-39 | 148 |
| **gene_233\|GeneMark.hmm\|350_aa\|+\|5216\|6268** | ammonia monooxygenase [Polymorphum gilvum] | gi\|503419284\|ref\|WP_013653945.1\| | 4.94E-67 | 223 |
| **gene_246\|GeneMark.hmm\|318_aa\|+\|6379\|7335** | ammonia monooxygenase [Azorhizobium caulinodans] | gi\|501122003\|ref\|WP_012171155.1\| | 1.60E-48 | 174 |
| **gene_196\|GeneMark.hmm\|336_aa\|-\|14681\|15691** | membrane protein [Pseudogulbenkiania sp. MAI-1] | gi\|635657271\|ref\|WP_024304138.1\| | 1.58E-85 | 270 |
| **gene_124\|GeneMark.hmm\|359_aa\|+\|83125\|84204** | hypothetical protein [Variovorax paradoxus] | gi\|951108200\|ref\|WP_057597630.1\| | 1.15E-103 | 317 |
| **gene_35\|GeneMark.hmm\|359_aa\|-\|30096\|31175** | hypothetical protein [Variovorax paradoxus] | gi\|951108200\|ref\|WP_057597630.1\| | 1.15E-103 | 317 |
